# Estimating Insulin Sensitivity and Beta-Cell Function from the Oral Glucose Tolerance Test: Validation of a new Insulin Sensitivity and Secretion (ISS) Model

**DOI:** 10.1101/2023.06.16.545377

**Authors:** Joon Ha, Stephanie T. Chung, Max Springer, Joon Young Kim, Phil Chen, Melanie G. Cree, Cecilia Diniz Behn, Anne E. Sumner, Silva Arslanian, Arthur S. Sherman

## Abstract

Efficient and accurate methods to estimate insulin sensitivity (S_I_) and beta-cell function (BCF) are of great importance for studying the pathogenesis and treatment effectiveness of type 2 diabetes. Many methods exist, ranging in input data and technical requirements. Oral glucose tolerance tests (OGTTs) are preferred because they are simpler and more physiological. However, current analytical methods for OGTT-derived S_I_ and BCF also range in complexity; the oral minimal models require mathematical expertise for deconvolution and fitting differential equations, and simple algebraic models (e.g., Matsuda index, insulinogenic index) may produce unphysiological values. We developed a new ISS (Insulin Secretion and Sensitivity) model for clinical research that provides precise and accurate estimates of SI and BCF from a standard OGTT, focusing on effectiveness, ease of implementation, and pragmatism. The model was developed by fitting a pair of differential equations to glucose and insulin without need of deconvolution or C-peptide data. The model is derived from a published model for longitudinal simulation of T2D progression that represents glucose-insulin homeostasis, including post-challenge suppression of hepatic glucose production and first- and second-phase insulin secretion. The ISS model was evaluated in three diverse cohorts including individuals at high risk of prediabetes (adult women with a wide range of BMI and adolescents with obesity). The new model had strong correlation with gold-standard estimates from intravenous glucose tolerance tests and hyperinsulinemic-euglycemic clamp. The ISS model has broad clinical applicability among diverse populations because it balances performance, fidelity, and complexity to provide a reliable phenotype of T2D risk.

## Introduction

Glucose homeostasis and risk of type 2 diabetes (T2D) depends chiefly on the balance of insulin sensitivity (S_I_) and pancreatic beta-cell function (BCF). Which factor predominates depends on the individual and the population (1–4). Efficient and effective methods for measuring the two factors are thus of great importance for early T2D risk stratification and prevention programs. We present here a novel method for assessing insulin sensitivity and beta-cell function using a standard oral glucose tolerance test (OGTT) with special consideration to data requirements, ease of implementation, and effectiveness. This model, the Insulin Sensitivity and Secretion (ISS) model, is designed to simultaneously estimate the two principal factors and give a complete picture of overall diabetes risk.

This innovative mathematical model is needed because currently available methods do not provide simultaneous estimates of insulin sensitivity and beta cell function using a minimally invasive protocol with judicious resource utilization (Supplemental Table 1; all Supplemental Material listed after the Discussion).

Current approaches range from highly complex to simplistic. At the high end of complexity and invasiveness are the intravenous methods for estimating insulin sensitivity and insulin secretion under defined, highly controlled conditions: the hyperinsulinemic-euglycemic clamp (HIEC), the hyperglycemic clamp (HGC), and the frequently sampled, insulin-modified intravenous glucose tolerance test (IM-FSIGT). The intravenous methods are reliable and reproducible and reflect maximal secretory responses to supra-physiologic or extreme hyperglycemic conditions but do not reflect postprandial physiologic responses. Most important, the intravenous methods are invasive, expensive, and require a high degree of technical expertise, increasing the burden on patients and study personnel.

Models that utilize oral glucose or meal perturbations to derive insulin sensitivity and beta cell function estimates address some of the limitations of intravenous methods (5). The oral minimal model (OMM) fits OGTT glucose concentration data to estimate insulin sensitivity, and the C-peptide minimal model fits OGTT C-peptide concentration data to estimate first- and second-phase beta-cell function. The rates of change of glucose and C-peptide are estimated from sparsely sampled data by deconvolution, which is technically challenging, and generally requires at least seven time points and durations longer than two hours during the OGTT (Supplemental Table S1). Ideally, models designed to leverage the more commonly performed two-hour OGTTs using sparsely collected timepoints (3–5) are practical and preferred. However, sometimes durations longer than two hours are required (5, 6).

The simplest equations for estimating diabetes risk are constructed from algebraic combinations of measurements taken in the fasting state or during OGTTs (for example, Matsuda insulin sensitivity index (ISI), insulinogenic index (IGI), homeostasis model assessment for insulin resistance and beta-cell function (HOMA-IR, HOMA-Beta; see Supplementary Table 1). These measures are widely used in clinical and epidemiologic studies because they do not require solution of differential equations or curve fitting and can be obtained from a single fasting blood draw or a five-point OGTT. However, algebraic indices are susceptible to unphysiological negative or outlier values especially when the number of time points are few or when fasting insulin is low or insulin secretion is impaired. The risk for unphysiological numbers is high in diverse populations where the contribution of insulin secretion and sensitivity to overall diabetes risk is heterogenous.

Given the challenges and limitations of the methods just described, we designed a novel pragmatic and versatile model for estimating insulin sensitivity and beta-cell function that uses insulin and glucose from a standard five-point OGTT. The goal of this work was to develop a model that balances performance with conservation of resources and patient burden. It can be deployed with reasonable accuracy and precision for assessing beta cell function and insulin sensitivity, along with glucose tolerance status, in large-scale clinical studies and across diverse populations. The ISS model requires solution and fitting of differential equations but does not require C-peptide data or the technical expertise of deconvolution. We apply the model to two independent, diverse data sets and show that the estimates strongly correlate with gold standard measures. We use these and an additional independent data set to show that acceptable estimates can also be obtained from three-point OGTTs. The modest data requirements and ease of use make it suitable for use in large-scale clinical studies and open up large collections of archival data for further analysis.

## Methods

To compare the performance of the ISS model to gold standard methods across age and ethnicity, we used two archival data sets, from The Federal Women’s Study (FWS) and The University of Pittsburgh Medical Center Children’s Hospital of Pittsburgh (UPMC CHP). To compare ISS model estimates derived from reduced sampling time points, we used three data sets: FWS, UPMC CHP, and an additional adolescent cohort from The University of Colorado (CU). Further details of the methods can be found in the Supplemental Methods.

### The Federal Women’s Study

The data for this analysis were obtained from women, 18-65 years old, in the Federal Women’s Study (ClinicalTrials.gov identifier, NCT01809288), who had an OGTT and IM-FSIGT, a trial designed to examine the racial/ethnic variations in the risk of diabetes and heart disease in African, African American, and white women (7–9). The demographic and metabolic characteristics of the 131 women from this study are provided in Table 1. Of these, 111 who had both OGTT and IM-FSIGT were used to compare estimates from the two tests (Figs. 1 – 4, S1), and 129 who had complete OGTTs were used to compare estimates with ISS using five time points (0, 30, 60, 90, 120 min) vs. three time points (0, 60, 90 min). (See Figs. 9, 11, S3).

**Figure 1.**
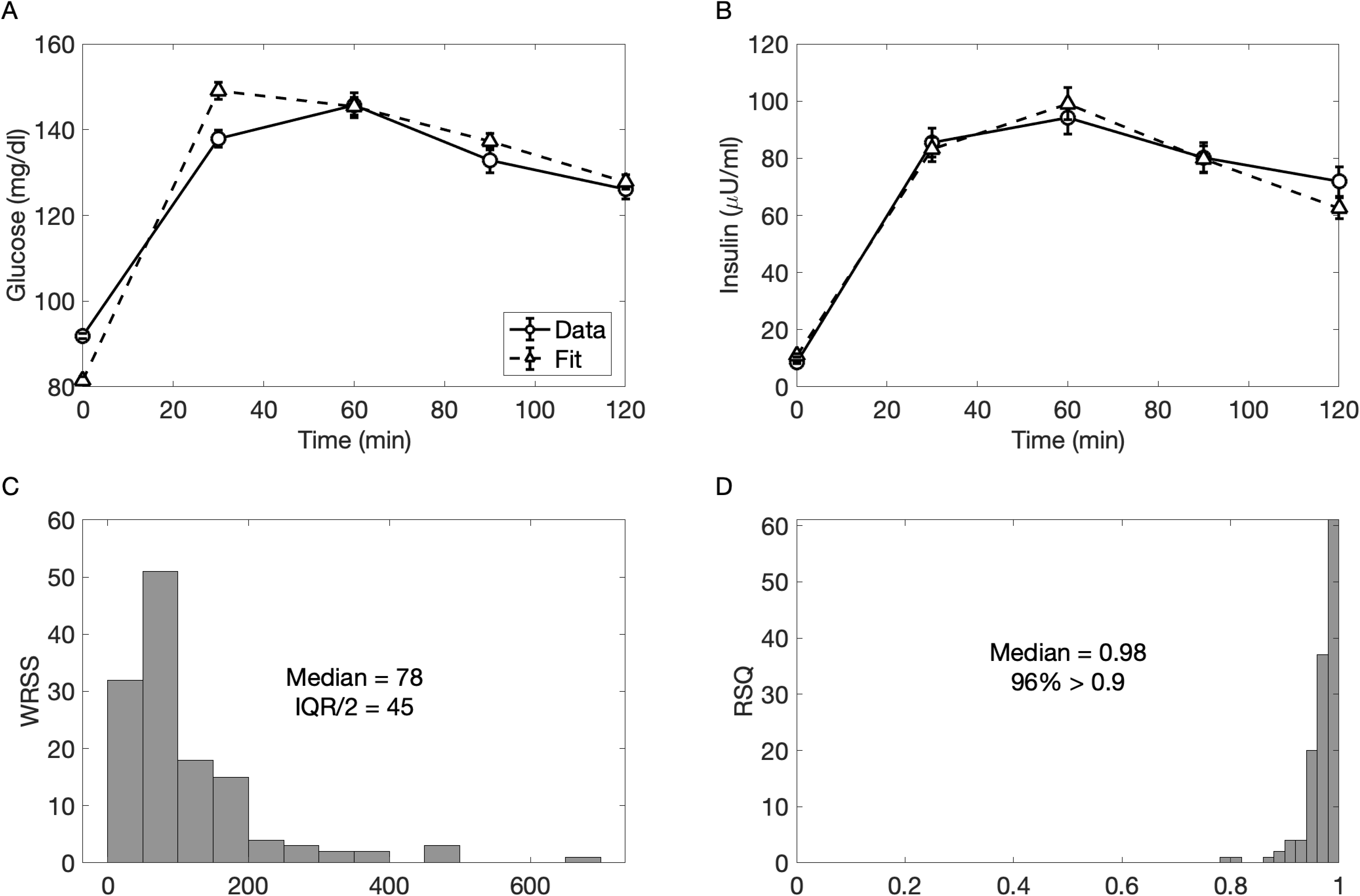
Fitting error for FWS. Data and fitted curves (mean ± SEM) for glucose (A) and insulin (B). Histogram of weighted residual sum of squares (WRSS) (C) and R^2^ (D).

**Table 1.** Characteristics of Federal Women’s Study Participants

|  | NGT<br>n=90 | PreDM<br>n=41 | P-value |
| --- | --- | --- | --- |
| Age (yrs.) | 40.6 ± 9.4 | 48.4 ± 8.4 | P < 0.001 |
| Race (%Black/White) | 59/41 | 54/46 | P = 0.7 |
| BMI (kg/m<sup>2</sup>) | 29.6 ± 5.9 | 30.7 ± 4.7 | P = 0.2 |
| HbA1c (%) | 5.3±0.4 | 5.6±0.3 | P < 0.001 |
| Fasting glucose<br>(mg/dL) | 89.4 ± 5.3 | 97.0 ± 7.6 | P < 0.001 |
| Fasting insulin<br>(μU/mL) | 16.9 ± 12.1 | 18.1 ± 11.1 | P = 0.6 |
| 2-hr glucose (mg/dL) | 113.4 ± 14.2 | 154.0 ± 21.8 | P < 0.001 |
| 2-hr insulin (μU/mL) | 47.0 ± 59.6 | 72.6 ± 98.9 | P = 0.003 |
| AIR<sub>g</sub><br>(μU/mL × min) | 861 ± 734 | 600 ± 380 | P = 0.01 |
| MinMod S<sub>I</sub><br>((μU/mL)<sup>-1</sup> min<sup>-1</sup> × 10<sup>4</sup>) | 3.17 ± 2.60 | 2.60 ± 1.57 | P = 0.01 |

Data are mean ± SD. NGT: normal glucose tolerance; PreDM: pre-diabetes.
Black includes African-American and African Immigrants; BMI: body mass index.
All participants were female.

During the IM-FSIGT, baseline blood samples were obtained, and dextrose (0.3 g/kg) was administered intravenously. Insulin was given as a bolus at 20 minutes (0.03 U/kg). As previously described, plasma glucose and insulin concentrations were obtained at 32 time points between baseline and 180 minutes (7). The acute insulin response to glucose (AIRg) was defined as the insulin area under the curve (AUC) greater than basal at 0 to 10 minutes (10). The insulin sensitivity index (S_I_) was calculated via the minimal model (MINMOD Millennium, version 6.02) (10).

### The University of Pittsburgh Medical Center Children’s Hospital of Pittsburg Adolescent Study (UPMC CHP)

The data of the 131 adolescents (age 10-20 years) from UPMC CHP study reported here were taken from two studies funded by the National Institutes of Health, Childhood Insulin Resistance and Childhood Metabolic Markers of Adult Morbidity in blacks (NCT00640224 and NCT01312051, respectively). The demographic characteristics of the participants are provided in Table 2. Information resulting from these grants unrelated to model fitting has been published previously (11–17).

**Table 2.** Characteristics of UPMC CHP Study Participants

|  | NGT (1)<br>n=68 | PreDM (2)<br>n=34 | T2D (3)<br>n=29 | P ANOVA | P, post hoc |  |  |
| --- | --- | --- | --- | --- | --- | --- | --- |
|  |  |  |  |  | 1 vs. 2 | 1 vs. 3 | 2 vs. 3 |
| Age (yrs.) | 14.9 ± 1.7 | 15.1 ± 1.8 | 15.1 ± 1.7 | NS | - | - | - |
| Race (% Black/White) | 44/56 | 35/65 | 52/48 | NS | - | - | - |
| Sex (% M/F) | 51/49 | 41/59 | 38/62 | NS | - | - | - |
| Tanner stage (% III/IV/V) | 15/12/73 | 9/12/79 | 0/28/72 | NS | NS | NS | NS |
| BMI (kg/m<sup>2</sup>) | 35.3 ± 5.7 | 37.2 ± 6.9 | 36.7 ± 5.3 | NS | - | - | - |
| BMI %ile | 98 ± 2 | 98 ± 1 | 99 ± 0.6 | NS | - | - | - |
| HbA1c (%) | 5.4 ± 0.3 | 5.4 ± 0.3 | 6.5 ± 0.8 | <0.0001 | NS | <0.0001 | <0.0001 |
| Fasting glucose (mg/dL) | 88.6 ± 6.2 | 92.7 ± 7.7 | 115.7 ± 27.9 | <0.0001 | 0.01 | <0.0001 | 0.0002 |
| Fasting insulin (μU/mL) | 28.9 ± 14.5 | 38.3 ± 22.7 | 64.5 ± 57.5 | <0.0001 | 0.03 | 0.002 | 0.03 |
| 2-hr glucose (mg/dL) | 114.8 ± 14.3 | 154.7 ± 14.5 | 202.4 ± 51.7 | <0.0001 | <0.0001 | <0.0001 | <0.0001 |
| 2-hr insulin (μU/mL) | 142.2 ± 116.9 | 311.8 ± 171.1 | 206.5 ± 173.6 | <0.0001 | <0.0001 | NS | 0.01 |
| Insulin sensitivity by clamp (mg/kg/min per μU/ml) | 2.4 ± 1.1 | 1.9 ± 1.2 | 1.3 ± 0.9 | <0.001 | NS | <0.001 | NS |
| 1<sup>st</sup>-phase insulin (μU/ml) | 246.4 ± 216.4 | 220.6 ± 123.3 | 124.7 ± 155.9 | 0.01 | NS | 0.01 | NS |
| 2<sup>nd</sup>-phase insulin (μU/ml) | 279.9 ± 168.3 | 326.1 ± 217.0 | 166.5 ± 133.4 | 0.001 | NS | 0.01 | 0.001 |

Data are mean ± SD. Ob: obese; NGT: normal glucose tolerance; IGT: impaired glucose tolerance; T2D: type 2 diabetes.
AA: African American; AW: American-White; BMI: body mass index

After a 10- to12-hr overnight fast, participants underwent a 2-hr OGTT (1.75 g/kg, maximum 75 g), and blood samples were obtained at −15, 0, 15, 30, 60, 90 and 120 min for the measurement of glucose and insulin. The day after the OGTT participants underwent either HIEC or HGC in random order 1 – 4 weeks apart (18). Each clamp evaluation was performed after a 10- to 12- hr overnight fast. In vivo insulin sensitivity was evaluated during a 3-hr hyperinsulinemic (80 mU/m^2^/min)-euglycemic clamp (100 mg/dL) (18). 1st- and 2nd -phase insulin secretion was assessed during a 2-hr hyperglycemic (225 mg/dL) clamp as described previously (18). Plasma glucose was increased rapidly to 225 mg/dL by a bolus infusion of dextrose and maintained at that level by a variable rate infusion of 20% dextrose for 2 hours, with frequent measurement of glucose and insulin concentrations.

During the hyperinsulinemic-euglycemic clamp, insulin-stimulated glucose disposal (Rd) was calculated to be equal to the rate of exogenous glucose infusion during the final 30 min of the clamp. Peripheral SI was calculated by dividing Rd by the steady-state clamp insulin concentration multiplied by 100 (18). During the HGC, beta cell function, measured as 1st- and 2nd -phase insulin concentrations, was calculated as previously described (11, 18).

### The University of Colorado Anschutz (CU) studies of adolescent women

Data were combined from five studies in female adolescents (12-21 years) with varied BMI and with or without polycystic ovary syndrome (PCOS). For this secondary analysis, only participants with a complete OSTT data set including all glucose and insulin values were included from the parent studies (N=113).

(19). In these studies, a modified oral sugar tolerance test (75 g glucose and 25 grams of fructose, OSTT) was performed with seven time points over 120 min (0, 10, 20, 30, 60, 90, 120). Because of the non-standard inclusion of fructose, the OSTT data were used only for the purpose of comparing estimates with different numbers of time points, not for comparing to gold-standard measures.

An additional data set of 33 adolescent females from the AIRS (Androgens and Insulin Resistance Study) study (20) was used that had 75-gram OGTTs, with blood draws at 0, 15, 30, 60, 90, 120 minutes, fit with the OMM using SAAM II (SAAM II software v 2.2, The Epsilon group, Charlottesville, VA, USA) to estimate S_I_, as well as a 4 phase hyperinsulinemic-euglycemic clamp. Phases included fasting, then consecutive insulin doses for each 1.5-hour phase of 10, 16, and 80 mU/m^2^/min. 20% dextrose was infused to maintain blood glucose at 95 mg/dL, with blood samples drawn every 5 minutes. GIR (mg/kg/min) or M-value, was determined based on steady-state measurements from the final 30 minutes of the 80 mU/m^2^/min phase of the clamp (21, 22).

### Glucose Tolerance Status

Across all three cohorts, glucose tolerance status was defined according to the American Diabetes Association guidelines. Prediabetes was defined as fasting glucose ≥ 100 but < 126 mg/dL and/or 2-hour glucose level ≥ 140 but < 200 mg/dL (29). Type 2 diabetes was determined based on documented history of chart review (UPMC) or established clinical presentation with type 2 diabetes and with HbA1c (UPMC and UC).

### Biochemical measurements

In FWS, glucose and insulin concentrations were measured in serum using the Roche Cobas 6000 analyzer (Roche Diagnostics, Indianapolis, IN). In the UPMC CHP cohort, plasma glucose was determined by the glucose oxidase method using a glucose analyzer (Yellow Springs Instrument Co., Yellow Springs, OH, USA), and insulin by commercially available radioimmunoassay (Millipore, St. Charles, MO, USA), as previously reported (15). HbA1c was measured by high-performance liquid chromatography (Tosoh Medics, Inc., San Francisco, CA, USA). In the CU study, blood glucose was measured using StatStrip ® Hospital Glucose Monitoring System (Novo Biomedical) except for AIRS (Yellow Springs Instrument Co., Yellow Springs, OH, US). Serum insulin was determined with radioimmunoassay (EMD Millipore, Billerica, MA), HbA1c was measured by high-performance liquid chromatography (Tosoh Medics, Inc., San Francisco, CA, USA) or with a handheld analyzer.

### Modeling methods

#### The model

The model for data fitting is adapted from our previously published model for simulating longitudinal progression to T2D (23, 24). The equations for glucose and insulin are:

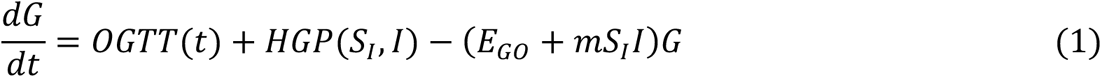

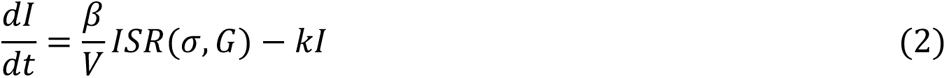

where *G* is glucose concentration (mg/dl), *I* is insulin concentration (μU/ml), and *OGTT*(*t*) is the glucose rate of appearance (*R*_*a*_) during the OGTT, modeled by a piecewise linear function similar to the approach in (25), as described in (24). Time is in min. *HGP*(*S_I_*, *I*) represents hepatic glucose production, which depends on peripheral insulin sensitivity (*S_I_*)—because peripheral and hepatic insulin sensitivity are correlated (24)—and also insulin concentration, which suppresses hepatic glucose production. Following the nomenclature in (26), *E_GO_* is the glucose effectiveness, representing the non-insulin dependent uptake of glucose, *β* is the beta-cell mass, *σ* is the beta-cell function (secretion per unit mass of beta cells), and *k* is rate of insulin clearance, primarily in the liver. *V* is the volume of distribution for insulin, and *ISR*(*σ*, *G*) is the insulin secretion rate. It is the output of a sub-model for exocytosis, which includes docked and readily releasable vesicles and simulates both first- and second-phase insulin secretion (27, 28). Details of parameters and functional forms are given in the Supplemental Equations. In general, model parameters, aside from the fitted ones described in the next two paragraphs, are the same as in (24), where all remaining details and rationales of the mathematical model can be found.

In (24) it was shown that the physiological detail included in the longitudinal model is sufficient to simulate the alterations in OGTTs as beta-cell function and mass slowly adapt in response to imposed reductions in *S_I_*, either leading to T2D or maintaining a non-diabetic state. Here the model is adapted for parameter estimation by fixing *β*, *σ*, and *S_I_*, which were slowly evolving variables in (24), making them parameters of the reduced system described by Eqs. 1 and 2 during OGTTs. The parameters *β* and *σ* appear as a product (because *ISR* in Eq. 2 is proportional to *σ*) and cannot be separately identified. Therefore, *β* is fixed at the value used in (24), and the parameters we estimate are *S*_*I*_ (mS_I_), and σ (mBCF).

#### Parameter Fitting Procedure

Estimation was done by fitting Equations (1) and (2) to the measured glucose and insulin concentrations during the OGTTs. The model parameters were determined using weighted least squares and the minimization function `fminsearch` in MATLAB (version 9.5.0 (R2021b), Natick, Massachusetts: The MathWorks Inc.). The weights were reciprocals of the standard deviations for each cohort calculated at each glucose and insulin time point. Bounds were imposed on the parameters to prevent them from becoming negative by setting the error to a large value if the fitting algorithm attempted a negative value. This was necessary because Eqs. 1, 2 become unstable, and *G* and *I* increase without bound if either *S*_*I*_ or σ becomes negative.

As an additional measure of goodness of fit, R^2^ was calculated as 1 – (sum squared error)/(sum squared data):

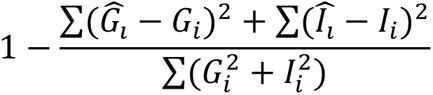

where the sums are over the time points of the OGTT, *G_i_* and *I_i_* are the measured glucose and insulin, and 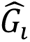 and 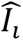 are the fitted values of glucose and insulin.

#### Determining Precision

Precision of the estimates was assessed by use of bootstrapping (29); because of the bounds on the parameter values simpler linearization methods are not valid. Samples were generated with replacement from the weighted residuals (WRSS values) for all individuals to create perturbed trajectories of G and I for each individual. We found 250 samples per individual to be adequate, as using 500 samples in a subset of cases did not appreciably change the results.

Coefficients of variation (CV) for each individual were assessed as the ratio of half the interquartile range (IQR/2) to the median of the bootstrap estimates.

#### Algebraic indices

The insulinogenic index (IGI), used to assess BCF, was calculated at the ratio of the incremental change of insulin and glucose from 0 to 30 minutes of the OGTT (14, 30, 31).

The Matsuda insulin sensitivity index (ISI) was calculated for two-hour OGTTs as (32)

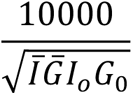

where *I*_0_ and *G*_0_ are the fasting insulin (μU/mL) and glucose (mg/dl) and 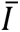 and 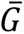 are the average insulin and glucose over the OGTT calculated as AUC/120 min by the trapezoidal rule.

Using the same definitions and units, HOMA IS, the reciprocal of HOMA IR, was calculated as (33):

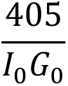

and HOMA Beta was calculated as (33):

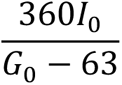

#### Measures of correlation and agreement

We used both Pearson and Spearman correlation coefficients and assessed agreement by Bland-Altman analysis (34). For the BCF measures, which were on widely different scales, we determined correlation coefficients only. To visually display agreement between *A* and *B*, where *A* and *B* were values obtained by two different methods or by the same method with different numbers of time points, we plotted *A* – *B* vs (*A* + *B*)/2, and to summarize agreement with a single number we calculated the relative pairwise difference Δ:

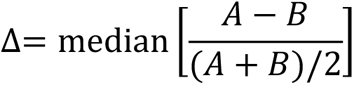

where the median was taken over the population. For comparison of differences in Δ between measures, we used the absolute value. For comparison with the literature, we also report pairwise percentage difference between *A* and reference value *B* as:

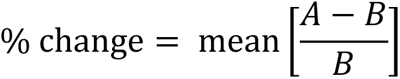

#### Statistics

We used Matlab (version 9.5.0 (R2021b) Natick, Massachusetts: The MathWorks Inc.), or R (R Foundation for Statistical Computing, ver. 4.1.1) for regression analyses. P < 0.05 was the criterion for statistical significance. To assess statistical significance of differences between non-independent quantities, such as correlations and relative differences (Δ) we used bootstrapping with 10,000 samples. Where dependence was not an issue, we used pairwise rank sum tests because of non-normality. NS is used to denote “not statistically significant”.

## Results

### The Federal Women’s Study and Comparison with Intravenous Glucose Tolerance Test

Figure 1 shows the mean fits for glucose (A) and insulin (B). Figure 1C shows a histogram of fitting error as weighted residual sum of squares (WRSS) for the cohort. The fits are good for most of the individuals albeit skewed, with a small fraction of outliers (9%) having error more than 1 SD above the mean (panel C). The median error was 78 and IQR/2 was 45. The histogram of values of R^2^ shows that 96% of the individuals had R^2^ > 0.9 (Figure 1D).

To assess the precision of the estimates, we carried out bootstrapping, as described in Methods. Figure 2 shows the histograms of estimates for mBCF (A) and mS_I_ (σ) (B) for a typical individual in the study. Figure 2C shows the scatter of the estimates in the mS_I_-σ plane. Boxplots for the CVs over all individuals are shown in Fig. 2D. The median CV was < 0.05 for both mS_I_ and 0, and CVs for mS_I_ were < 0.10 for 98% of the individuals and for σ were < 0.10 for 95% of the individuals. In summary, the precision was tight for almost all individuals except for a small subset of outliers.

**Figure 2.**
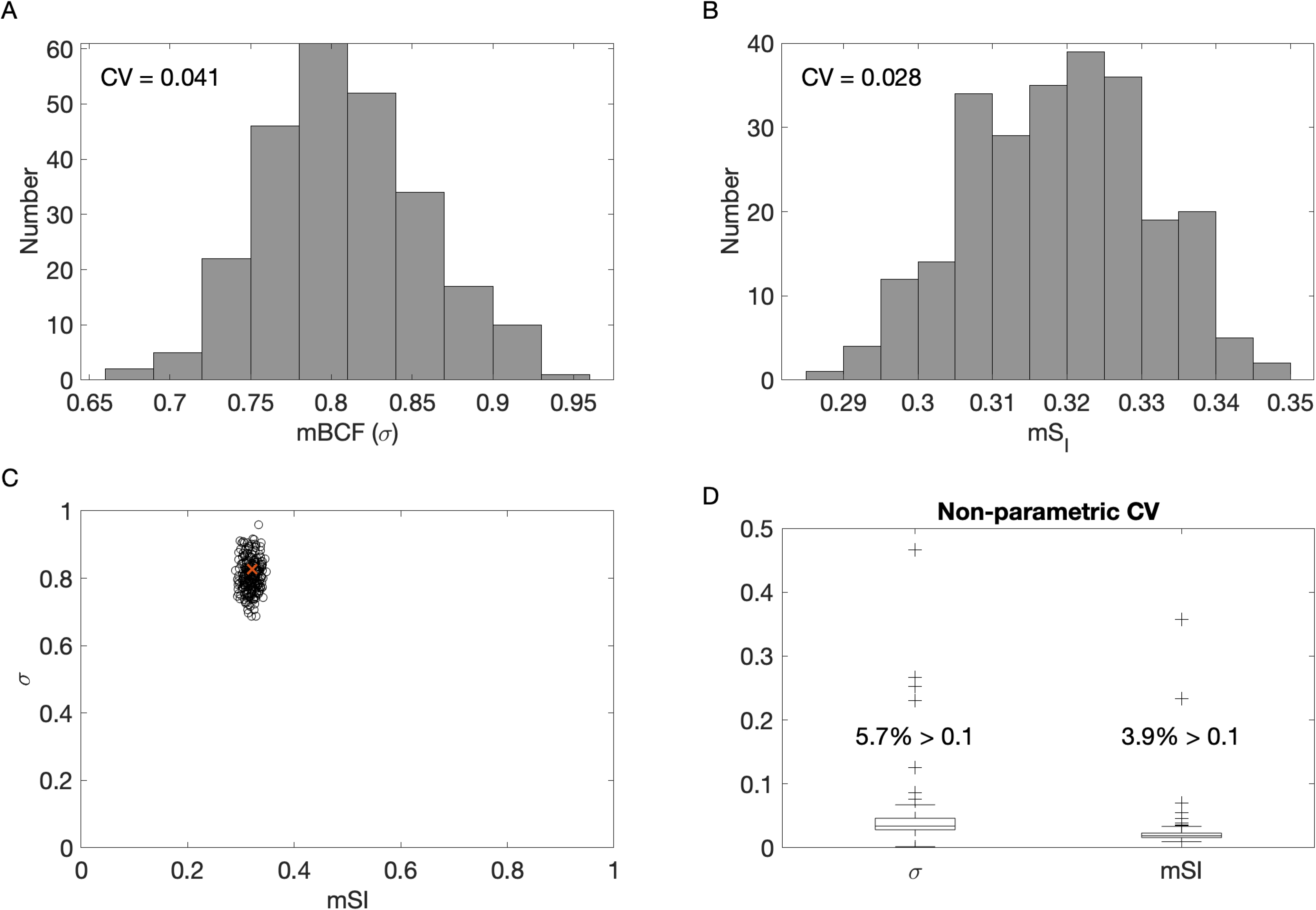
Precision of estimates for FWS by bootstrapping. Histogram of estimates for mBCF (σ) (A) and mS_I_ (B) and scatter plot of estimates in the mS_I_-σ plane (C). CV (Coefficient of Variation) is non-parametric (see Methods). Red X in C denotes the original fitted value used for analyses before perturbing with resampled residuals. D: box and whisker plots of the CVs for sigma and SI. Three outliers above 0.5 have been removed (0.66 and 0.99 for σ; 1.8 for mS_I_).

In most cases there was correlation between the bootstrap estimates for mS_I_ and σ (median = 0.45, IQR/2 = 0.31) indicating that there may be some compensation between these estimated parameters. However, despite this correlation, there were tight bounds and precise estimates for both parameters.

We next checked correlation and agreement with the S_I_ reported by MINMOD applied to IM-FSIGT data. There was moderate-strong correlation (Fig. 3A), and good agreement (Fig 3B) between mS_I_ and MINMOD S_I_. (See Table 4 for Pearson and Spearman correlation coefficients and median relative difference, Δ.) The Bland-Altman plot (Fig. 3B) shows a small bias (< 10%) and no significant slope for the regression of difference vs. the mean.

**Figure 3.**
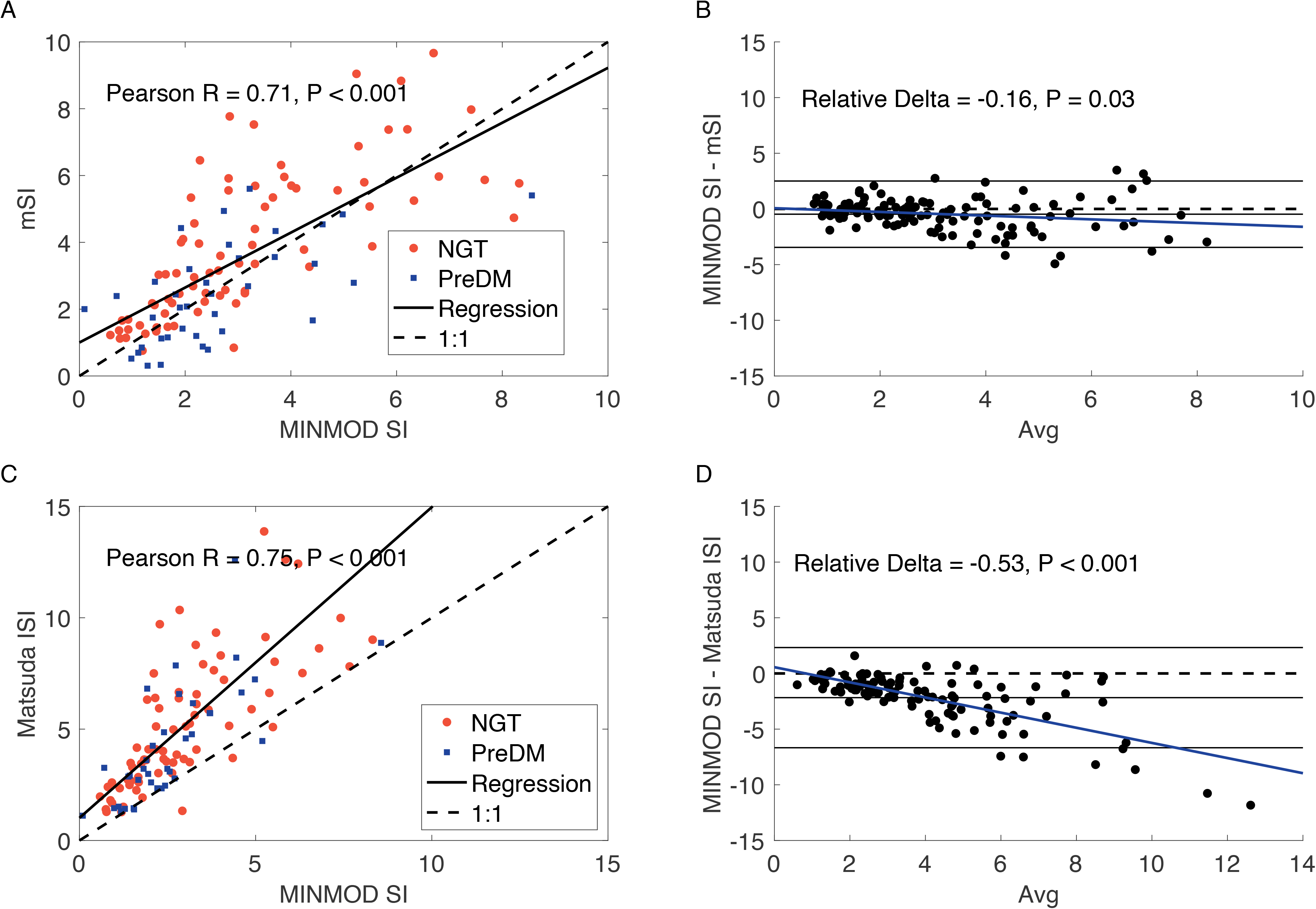
FWS insulin sensitivity measures. Correlation (A) and agreement assessed by Bland-Altman (B) for the ISS model mS_I_ compared to MINMOD S_I_. Correlation (C) and agreement assessed by Bland-Altman (D) for the Matsuda index compared to MINMOD SI. P values in C and D are for regression of difference vs. avg.

**Table 3.** Characteristics for UC Anschutz Study who completed OSTT

|  | NGT<br>n=84 | PreDM<br>n=29 | *P* t-test |
| --- | --- | --- | --- |
| Age (yrs.) | 15.9 ± 1.7 | 15.6 ± 1.6 | NS |
| Race (%White/Black/other) | 83/7/10 | 79/3/17 | NS |
| Sex (% M/F) | 0/100 | 0/100 | NS |
| Tanner stage (% III/IV/V) | 0/0/100 | 0/0/100 | NS |
| BMI (kg/m<sup>2</sup>) | 35.3 ± 5.9 | 37.5 ± 6.6 | NS |
| BMI %ile | 97.4 ± 2.1 | 98.3 ± 1.4 | NS |
| HbA1c (%) | 5.3 ± 0.2 | 5.8 ± 0.1 | <0.001 |
| Fasting glucose (mg/dL) | 89.5 ± 8.1 | 93.2 ± 8.4 | 0.04 |
| Fasting insulin (μU/mL) | 20.5 (15.0, 28.3) | 25.0 (14, 26) | NS |
| 2-hr glucose (mg/dL) | 132.7± 23.3 | 144.6 ± 28.6 | 0.05 |
| 2-hr insulin (μU/mL) | 158 (87.5, 206) | 314 (160, 600) | NS |

OSTT: oral sugar tolerance test (see Methods); mean±sd or median (interquartile range).

**Table 4.** Comparison of measures from this study with clamp and IM-FSIGT

| Insulin Sensitivity |  |  |  |  |  |
| --- | --- | --- | --- | --- | --- |
| Measure |  | Correlation and agreement with Hyperinsulinemic Euglycemic Clamp | Figure | Correlation and agreement with IM-FSIGT MINMOD S<sub>i</sub> | Figure |
| mS<sub>i</sub> |  | Pearson R = 0.66<br>Spearman Rho = 0.73<br>Δ = 0.49 | 7A | Pearson R = 0.71<br>Spearman Rho = 0.76<br>Δ = -0.16 | 3A |
| Matsuda ISI |  | Pearson R = 0.75***<br>Spearman Rho = 0.81***<br>Δ = 0.30*** | 7C | Pearson R = 0.75<br>Spearman Rho = 0.82**<br>Δ = -0.53*** | 3B |
| HOMA IS |  | Pearson R = 0.66<br>Spearman Rho = 0.78<br>Δ = 1.70*** | S1C | Pearson R = 0.71<br>Spearman Rho = 0.76<br>Δ = 1.20*** | S1A |
| Beta cell function |  |  |  |  |  |
| Measure |  | Correlation with Hyperglycemic Clamp | Figure | Correlation with AIRg | Figure |
| mBCF (s) | 1<sup>st</sup> phase | Pearson R = 0.70<br>Spearman Rho = 0.79 | 8A | Pearson R = 0.68<br>Spearman Rho = 0.70 | 4A |
|  | 2<sup>nd</sup> phase | Pearson R = 0.64<br>Spearman Rho = 0.78 | 8B |  |  |
| AUC I/AUC G | 1<sup>st</sup> phase | Pearson R = 0.62<br>Spearman Rho = 0.69 | 8C | Pearson R = 0.67<br>Spearman Rho = 0.68 | 4B |
|  | 2<sup>nd</sup> phase | Pearson R = 0.72<br>Spearman Rho = 0.83 | 8D |  |  |
| HOMA-Beta | 1<sup>st</sup> phase | Pearson R = 0.45**<br>Spearman Rho = 0.56** | S2C | Pearson R = 0.57<br>Spearman Rho = 0.62 | 4C |
|  | 2<sup>nd</sup> phase | Pearson R = 0.49*<br>Spearman Rho = 0.66** | S2D |  |  |
| IGI | 1<sup>st</sup> phase | Pearson R = 0.71<br>Spearman Rho = 0.79 | S2A | Pearson R = 0.52<br>Spearman Rho = 0.73 | 4D |
|  | 2<sup>nd</sup> phase | Pearson R = 0.63<br>Spearman Rho = 0.65 | S2B |  |  |

mBCF (s): beta-cell function from our model. Δ: agreement measured as median relative difference, defined in Methods. P < 0.001 for all correlations with clamp and IM-FSIGT. P values vs. corresponding value of mS<sub>i</sub> or s: , < 0.1; \*, < 0.05; \*\*, < 0.01; \*\*\*, < 0.001; no symbol: NS.

We also compared the relationships of established algebraic indices Matsuda ISI (Figs. 3C, D) and HOMA IS with MINMOD S_I_, (Fig S1A, B). Both Matsuda ISI and HOMA IS had moderate-strong correlation with MINMOD S_I_, but poor agreement. Matsuda ISI systematically overestimated S_I_ and HOMA IS systematically underestimated S_I_, evidenced by the regression slopes far from 1 and confirmed by the Bland-Altman plots (Figs. 3D, S1B). The median relative difference between MINMOD S_I_ and ISI was 4.6 times larger in magnitude than between MINMOD S_I_ and. mS_I_ and for HOMA IS was 7.5 times larger (P < 0.0001 by bootstrap for both).

Given that both mS_i_ and Matsuda ISI correlated strongly with MINMOD S_I_, it is not surprising that they correlated strongly with each other (0.9 for both Pearson R and Spearman Rho, P < 0.001).

We next compared the measure for BCF from our model with AIRg from IM-FSIGT. Our mBCF (σ) correlated moderately with AIRg (Fig. 4A). We also show for comparison the correlation between other commonly used measures of BCF based on OGTTs and AIR: AUC I/AUC G (Fig. 4B), HOMA Beta (Fig. 4C), and IGI (Fig. 4D). The correlation for mBCF with AIRg was numerically larger than each of those, but the differences were not statistically significant (Table 4).

**Figure 4.**
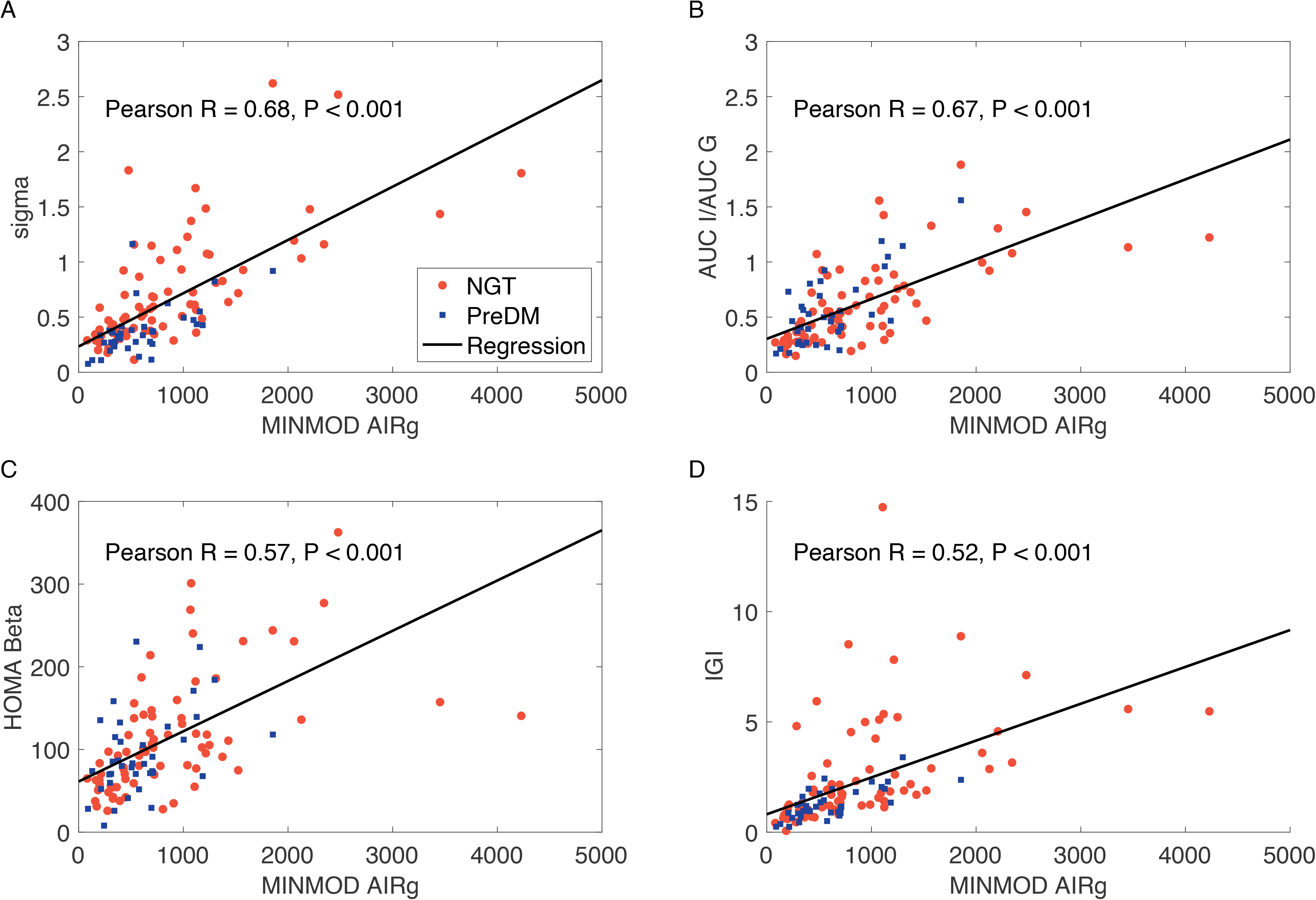
FWS BCF measures. BCF assessed by the ISS model σ correlated with AIRg from IM-FSIGT (A); Spearman rho = 0.70. Algebraic indices AUC I/AUC G (B), HOMA Beta (C) and insulinogenic index (D) compared to AIRg.

### The UPMC CHP Adolescent Study and Comparison with Clamp Studies

Figure 5 shows the mean fits for glucose (A) and insulin (B) and the WRSS error histogram (C). The histogram of values of R^2^ shows that 82% of the individuals had R^2^ > 0.9 (Figure 5D). The general trend of the data is captured well, but is skewed, similar to FWS (Figure 1). The error in the UPMC CHP fits is larger and more spread out than for FWS, with a median of 123 and IQR/2 of 97. The insulin levels were more than twice as high as for FWS, as expected for adolescents with severe insulin resistance.

**Figure 5.**
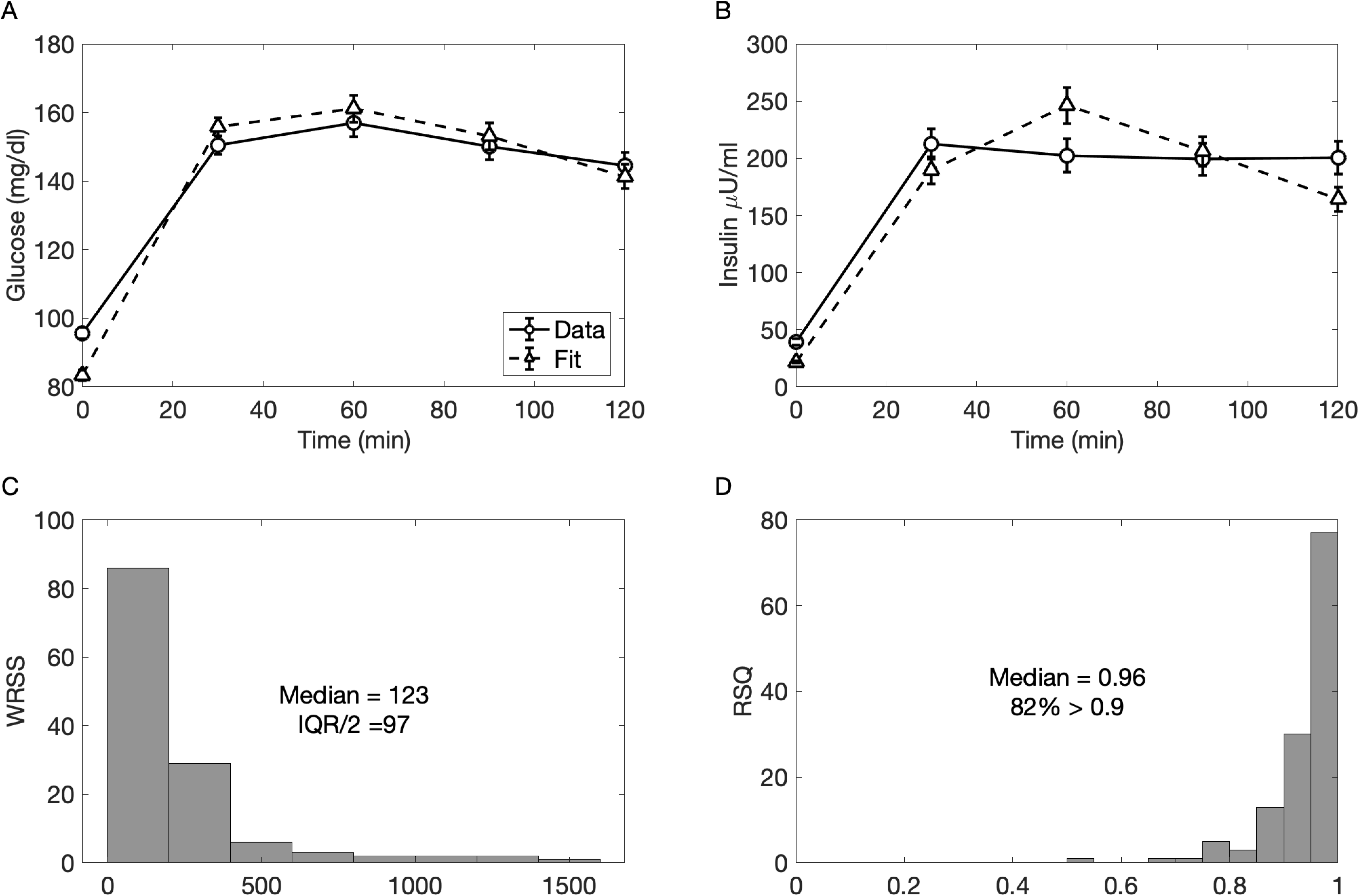
Fitting error for UPMC CHP study. Data and fitted curves (mean ± SEM) for glucose (A) and insulin (B). Histogram of weighted residual sum of squares (WRSS) (C) and R^2^ (D).

Figure 6 shows the histograms of the bootstrap estimates for mBCF(A) and mS_I_ (σ) (B) for a typical individual in the study. The scatter of the estimates in the mS_I_-σ plane is shown in Fig. 6C. Boxplots for the CVs over all individuals are shown in Fig. 6D. The median CV was < 0.05 for both mS_I_ and mBCF (σ). CVs for mS_I_ were < 0.10 for 97% of the individuals and for mBCF (σ) were < 0.10 for 94% of the individuals. As for the FWS study, the precision was good for almost all individuals except a small subset of outliers, and the estimates were correlated (median R = 0.46, IQR/2 = 0.29).

**Figure 6.**
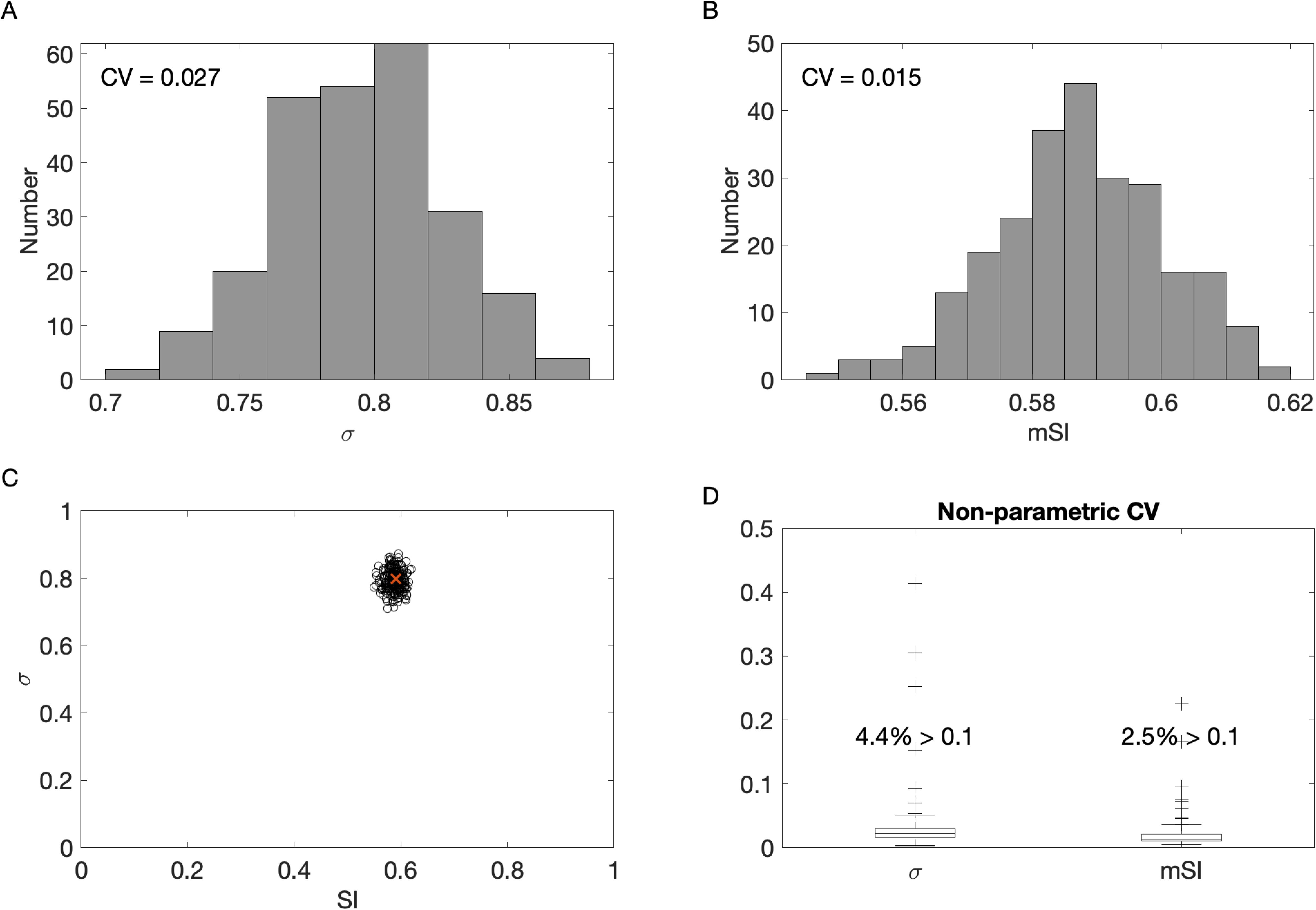
Precision of estimates for UPMC CHP study by bootstrapping. Histogram of estimates for mBCF (σ) (A) and mS_I_ (B) and scatter plot of estimates in the mS_I_-σ plane (C). CV (Coefficient of Variation) is non-parametric. Red X in C denotes the original fitted value used for analyses before perturbing with resampled residuals. D: box and whisker plots of CVs for sigma and SI. Four outliers with CV above 0.5 removed (range 0.54 – 0.61).

Figure 7 shows the correlation (7A) and agreement (7B) between mS_I_ and clamp SI (cS_I_). The correlation is comparable to that between mSI and MINMOD S_I_, but mSI was systematicallylower than cS_I_, as evidenced by the slope < −1 in 7A and the Bland-Altman plot in 7B. The error and variance were also not uniform, increasing roughly proportionally with cS_I_.; plotting logs makes the variance uniform (Fig.7A, inset). Figs. 7 C, D show that the Matsuda index similarly had good correlation with cS_I_ and underestimated cS_I_. Matsuda and mS_I_ were strongly correlated and in good agreement (not shown). Figures S1C, D show moderate correlation between HOMA IS and cS_I_ but, poor agreement with HOMA IS values much lower than cS_I_. See the Discussion for further details and comparison to other models.

**Figure 7.**
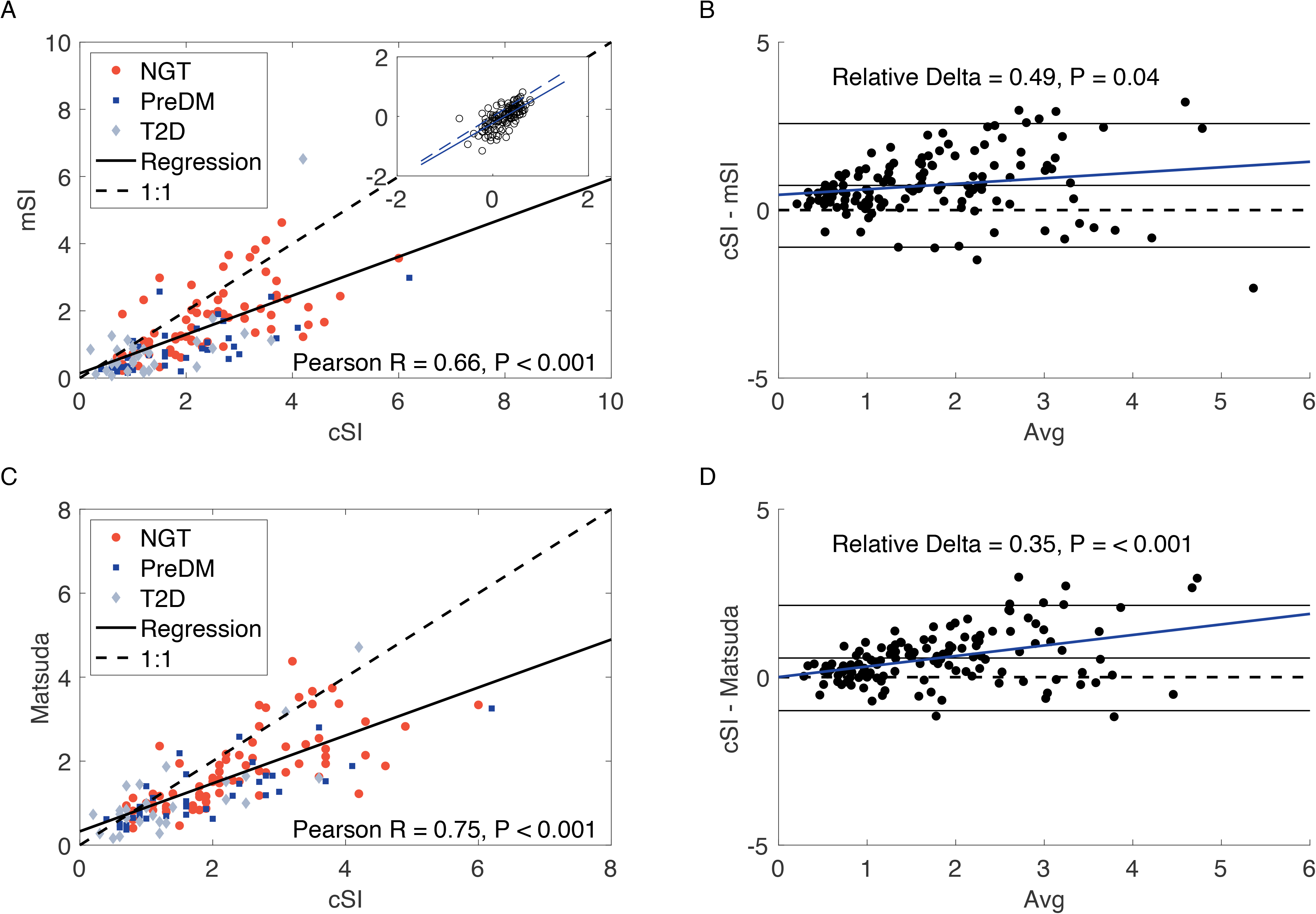
UPMC CHP SI Measures. Correlation (A) and agreement assessed by Bland-Altman (B) for our model mS_I_ compared to insulin sensitivity measured by hyperinsulinemic euglycemic clamp (cS_I_). mS_I_ underestimated cS_I_ pairwise by 31% (mean±SEM: 1.34±0.09 vs. 2.08±0.08, P < 0.001 by ranksum test). Correlation (C) and agreement assessed by Bland-Altman (D) for the Matsuda index compared to cS_I_. Matsuda underestimated cS_I_ pairwise by 20% (mean±SEM: 1.52±0.98 vs. 2.08±0.08, P < 0.001 by rank sum test). P values in C and D are for regression of difference vs. avg.

Figure 8 shows the moderate-strong correlation between mBCF (σ) and insulin secretion from HGC (first phase, panel A; second phase, panel B). The correlation of AUC I/AUC G with clamp HGC did not differ statistically from that of mBCF (Table 4). The correlations of insulinogenic index (IGI) with clamp first- and second-phase insulin secretion also did not differ statistically from those of mBCF (Figs. S2 A, B), but we note that one participant had a negative IGI because insulin at 30 min was lower than at time 0. The correlation of HOMA Beta with clamp HGC was about one-third smaller than that of mBCF (Figs. S2 C, D, Table 4).

**Figure 8.**
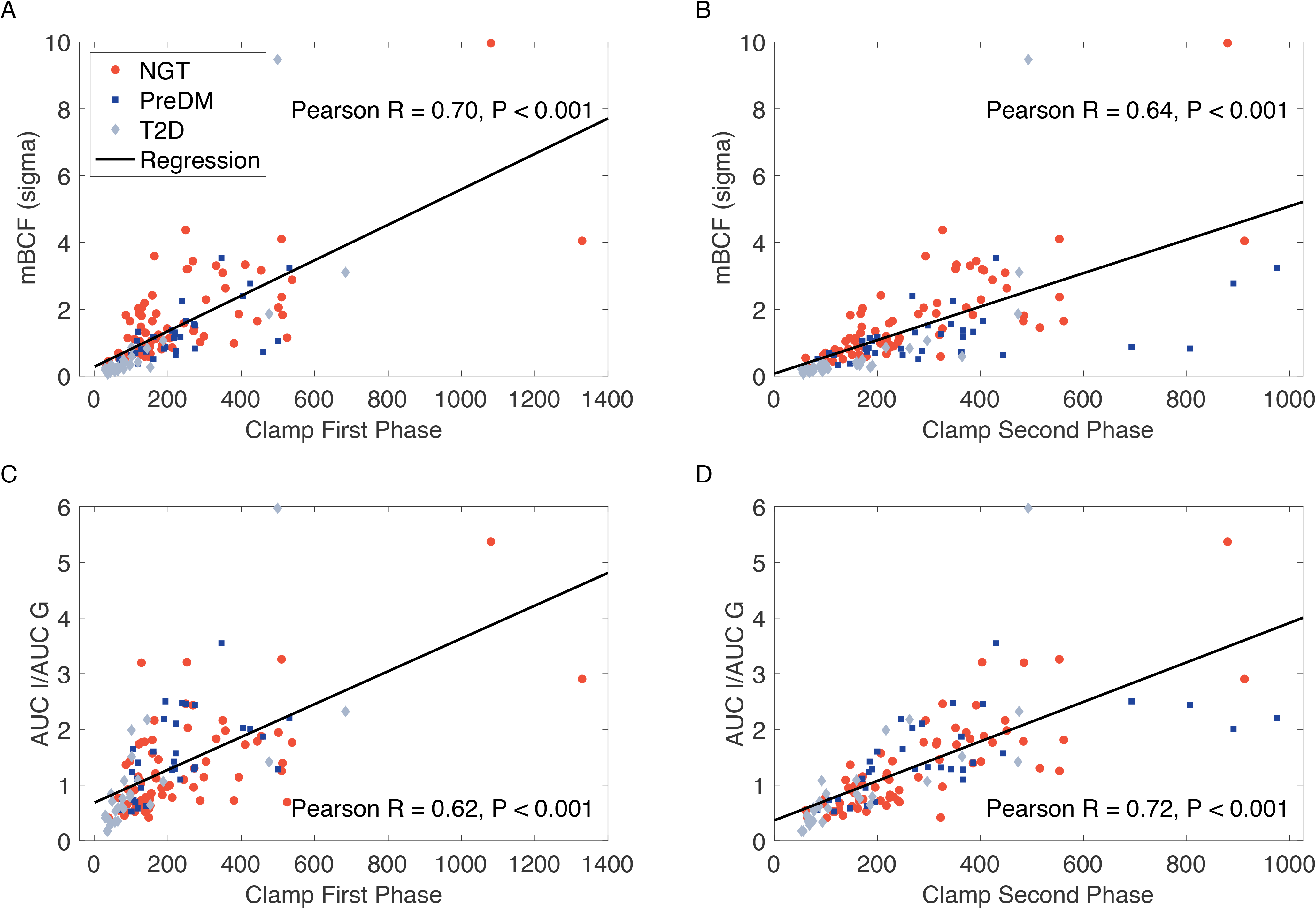
UPMC CHP BCF Measures. BCF assessed by our model σ correlated with first-phase (A) and second-phase insulin (B) from hyperglycemic clamp (HGC). BCF assessed by AUC I/AUC G correlated with first-phase (A) and second-phase insulin (B) from hyperglycemic clamp (HGC). (For comparison of HOMA Beta and insulinogenic index with HGC see Fig. S2.)

### ISS model estimates with reduced sampling time points (FWS and UPMC CHP)

Fig. 9 shows correlation and agreement for mS_I_ and mBCF (σ) using five time points (0, 30, 60, 90, 120) vs three time points (0, 60, 90) in the FWS study. For a fair comparison, WRSS error was calculated at all five time points (Fig. S3; statistics and details in figure legend). Parameter estimates, however, were done using five or three time points. The estimates for mSI with five and three time points were strongly correlated (R = 0.93, Fig. 9A) and showed no significant disagreement by Bland-Altman analysis (Fig. 9B) or pairwise comparison. The estimates for mBCF (σ) with five and three time points were also strongly correlated (R = 0.92, Fig. 9C) but the values using only three time points were systematically lower by Bland-Altman analysis (Fig. 9D). Pairwise, σ with three points was 21% lower than with 5 points.

**Figure 9.**
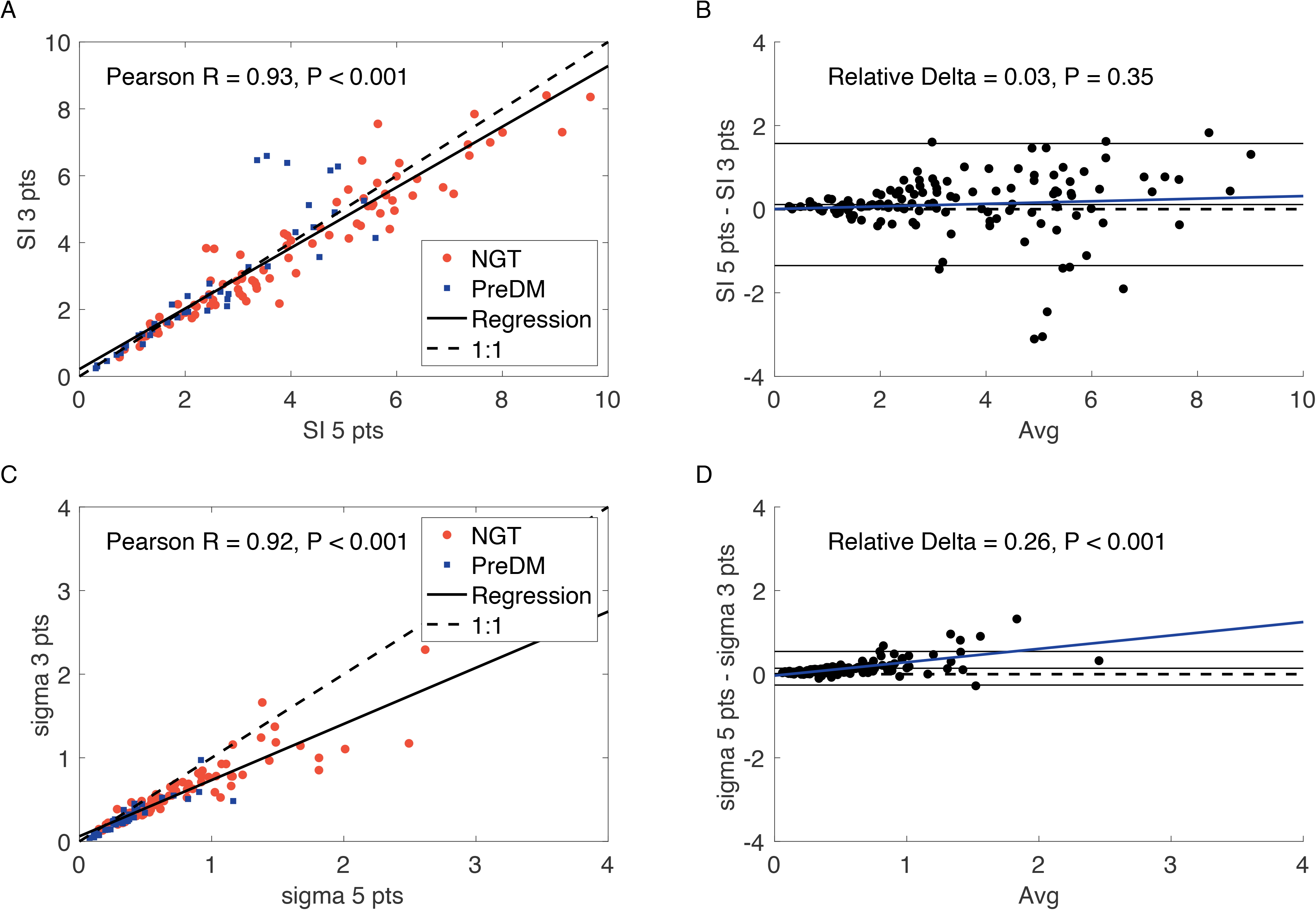
Reduced data sampling (3 points vs 5 points) for FWS. Comparison of parameter estimates with 5-point (t = 0, 30, 60, 90, 120 min) and 3-point OGTTs (t = 0, 60, 120 min) for FWS. Correlation (A) and agreement assessed by Bland-Altman analysis (B) for mS_I_. mS_I_ with 3 points was pairwise 2% lower (NS) than with 5 points (mean±SEM: 3.39±0.18 vs 3.5±0.16, P = 0.67 by ρανκσυμ test). Spearman Rho = 0.95. Correlation (C) and agreement assessed by Bland-Altman analysis (D) for mBCF (σ). Spearman Rho = 0.96. σ with 3 points was pairwise 21% lower than with five points (mean±SEM: 0.48 ±0.03 vs. 0.62±0.041, P = 0.007 by rank sum test). P values in C and D are for regression of difference vs. avg.

The same comparisons for the UPMC CHP study are shown in Fig. 10. Median WRSS increased from 123 to 160 when only three time points (t = 0, 60, 120 min) were used (Fig. S4). The correlation of the mS_I_ estimates was similar to that for FWS, (R = 0.88, Fig. 10A), and the values using only three time points were pairwise only 7% higher (NS), but the Bland-Altman regression was significant because the discrepancy was not uniform (Fig. 10B). For mBCF (σ), the correlation was lower than for FWS (Pearson R = 0.73, Fig. 10C); it rises to 0.80 if the two high leverage points with σ > 9 are excluded. The values for σ were 11% lower pairwise with three time points than with five, and the Bland-Altman regression was again significant because of non-uniform discrepancy (Fig. 10D).

**Figure 10.**
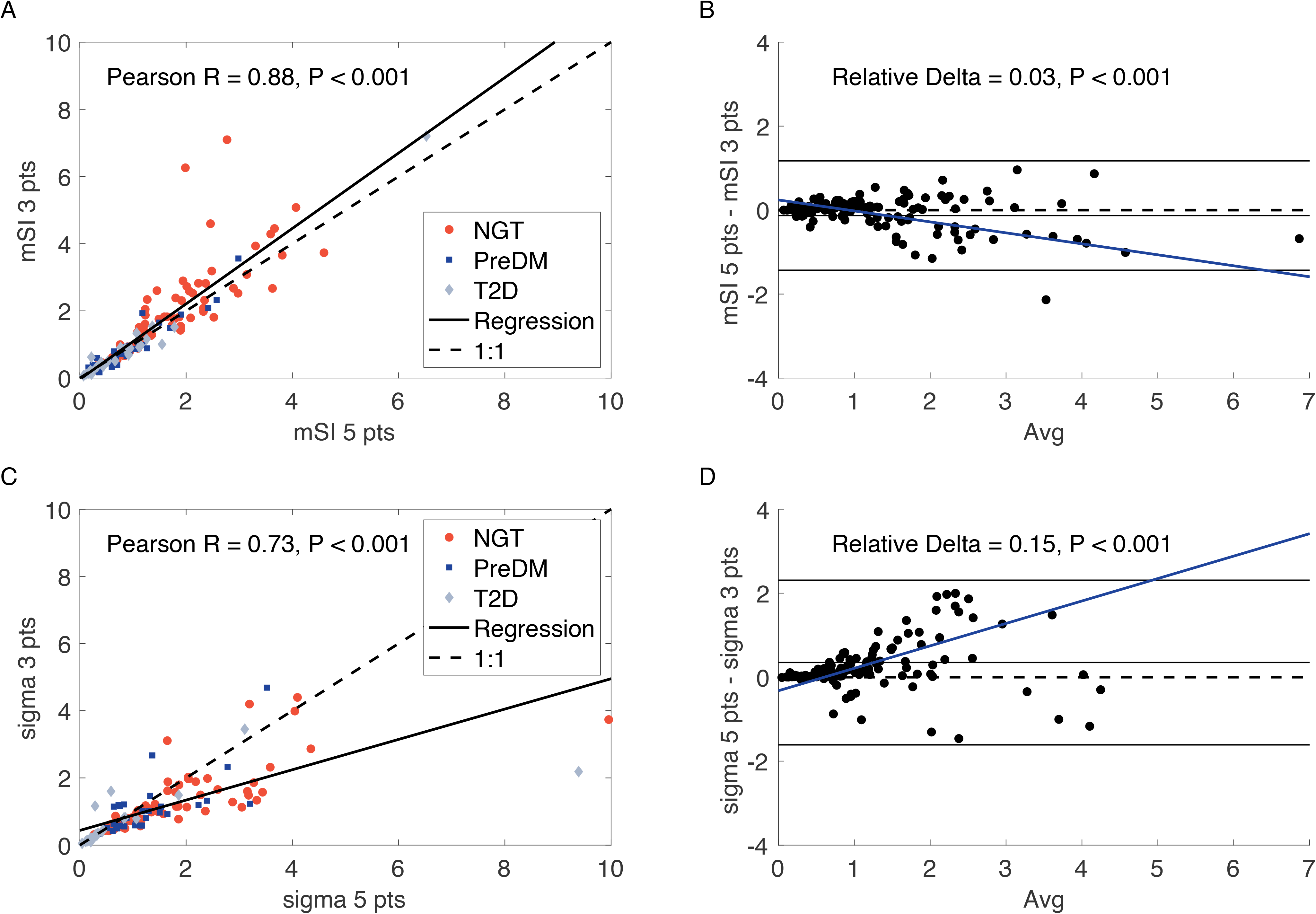
Reduced data sampling (5 points vs 3 points) for UPMC CHP. Comparison of parameter estimates with 5-point (t = 0, 30, 60, 90, 120 min) and 3-point OGTTs (t = 0, 60, 120 min) for UPMC CHP study. Correlation (A) and agreement assessed by Bland-Altman analysis (B) for model SI. mS_I_ was pairwise 7% higher (NS) with 3 time points (mean±sem: 1.49±0.12 vs. 1.36±0.09, P = 0.92 by paired ranksum test). Correlation (C) and agreement assessed by Bland-Altman analysis (D) for model sigma. If the two high-leverage points in C with s > 9 are excluded, Pearson R = 0.80. Two points with values > 4 (6.2, 7.2) clipped for display only in (D). Spearman Rho = 0.95 (A), 0.88 (C). σ was pairwise 11% lower with 3 time points (mean±sem: 1.08± 0.08 vs. 1.43±0.13, P = 0.02 by paired rank sum test). P values in C and D are for regression of difference vs. avg.

### ISS model estimates with reduced sampling time points (CU)

The Oral Glucose and C-peptide Minimal Models for insulin sensitivity and beta-cell function generally require at least 7 time points (5). We therefore used the UC data set, which had seven time points between 0 and 120 min (20) to test whether the ISS model would perform better with seven points than with five. Figs. S5 and S6 show fitting and error with five and seven time points, respectively, with the error calculated at the common five points. Adding two time points at t = 10 and 20 min, as done in the minimal models, did not improve the fit at the standard five points: WRSS ± IQR/2 was 150.8 ± 96.3 at the five points when five points were fit and 166.1 ±115.3 when seven points were fit. Also, the estimates of mS_I_ and σ using five points and seven points were very highly correlated with no significant systematic disagreement (Fig. S7). Finally, we used the UC data set to compare fitting with five time points vs three, and the results were similar to what we found with the FWS and UPMC CHP data sets (Fig. S8).

As a further demonstration of the acceptability of using three time points to fit the ISS model, Figure 11 shows for each of the three data sets that the model estimates of S_I_ and σ are effective at stratifying the participants by glucose tolerance status whether 5 points or 3 points are used. In all cases, the estimate pairs are shifted progressively towards the origin (low S_I_, low σ) as glucose tolerance worsens. Nonetheless, the two adolescent populations (panels C– F) tend to have lower S_I_ and higher σ than the adult population (panels A, B), as expected (35).

**Figure 11.**
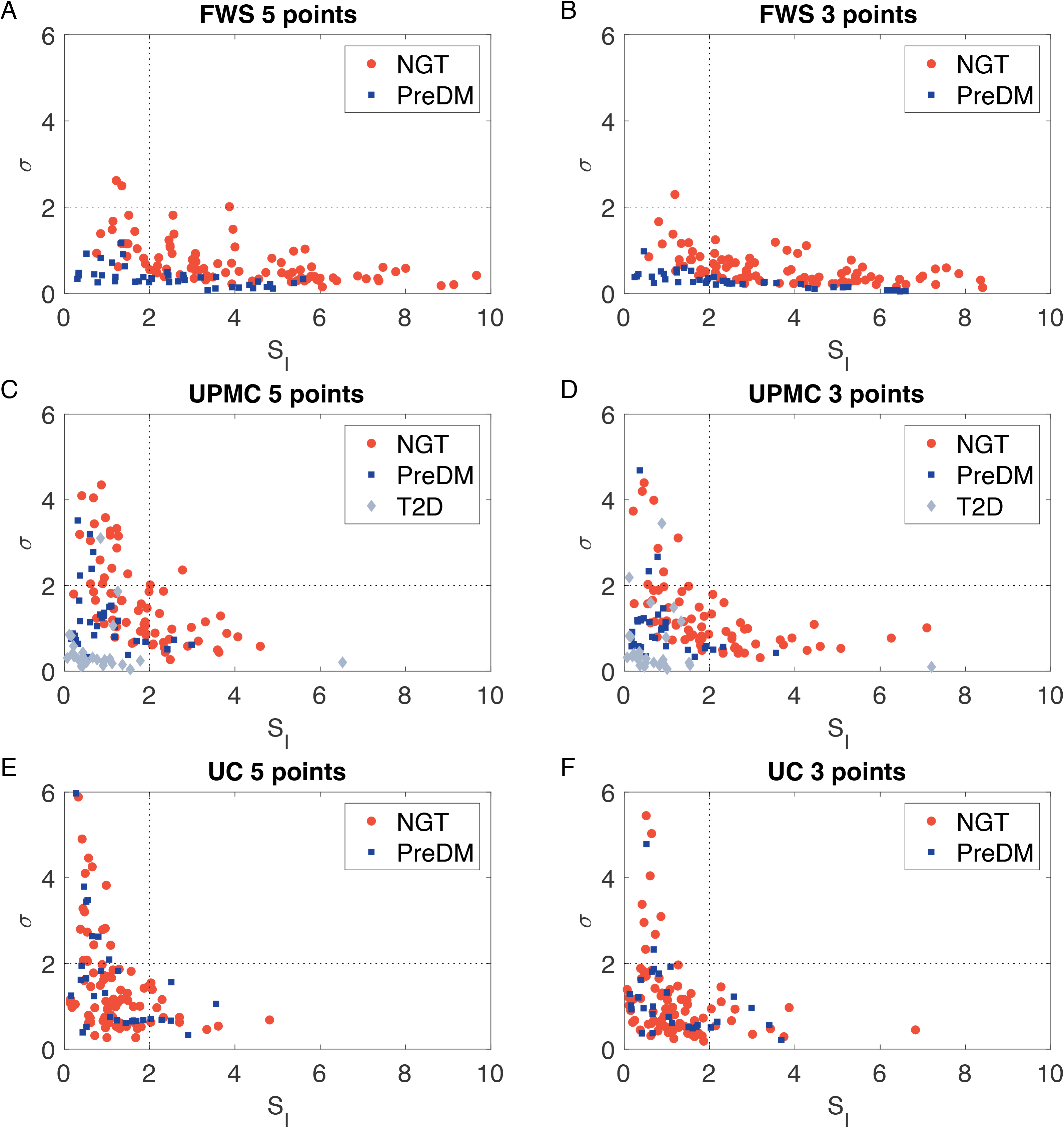
Stratification by glucose tolerance status with 5 or 3 time points. S_I_-σ pairs are plotted for the FWS, UPMC CHP and CU data sets. The dotted lines at S_I_ = 2 and s = 2 roughly divide the plane into quadrants of distinct glucose tolerance, with poorer tolerance points clustering in the lower right quadrant.

## Discussion

We introduced a new method for estimating insulin sensitivity and beta-cell function by fitting a system of ordinary differential equations describing glucose and insulin dynamics during a standard five-point OGTT. The model has high precision and good fits in two diverse cohorts across a wide range of age and BMI. Coefficient of variation was less than 10% for more than 95% of the participants studied. To assess accuracy, we compared the method with standard but more burdensome methods using IM-FSIGT and clamp (HIEC and HGC). We carried out the comparison using two cohorts with a wide age and BMI-range, the Federal Women’s Study (IM-FSIGT analyzed with MINMOD) and a study of adolescents with obesity conducted at the University of Pittsburgh Medical Center Children’s Hospital (HIEC and HGC). We found that our model estimates for insulin sensitivity (mS_I_) strongly correlated and had good agreement with two widely used measures, S_I_ obtained by fitting IM-FSIGT with MINMOD and peripheral insulin sensitivity obtained from the HIEC. The estimates from those cohorts for beta-cell function (mBCF) correlated well with those from IM-FSIGT (AIR) and HGC (both first and second phase).

### Comparison with algebraic indices

We also compared estimates of insulin sensitivity and beta cell function from the ISS model with commonly used algebraic indices for OGTTs (Matsuda ISI and HOMA IS for insulin sensitivity, and the insulinogenic index, HOMA beta, and AUC G/AUC I for beta-cell function). The correlation and agreement of the ISS model parameters and these algebraic indices with the standard measures are summarized in Table 4. Despite differences in agreement that reflect differences in the calculated measures, the ISS model parameters were highly correlated with the algebraic indices overall. We found that the ISS model avoids some deficiencies found with the algebraic indices, as detailed below. However, in comparison to other differential equation-based fitting methods, the ISS model obtains these improvements with modest requirements for computational effort and expertise associated with data fitting. It is possible to fit data from a medium-sized clinical study of several hundred participants in 5 – 10 minutes using a typical laptop or desktop computer. The Matlab scripts for our method are freely available (doi: 10.6084/m9.figshare.23535612) to make it as easy as possible for clinical researchers to employ it.

The ISS model mSI is preferable to Matsuda ISI in a key respect: the Matsuda index is sensitive to the values of fasting glucose and insulin, which appear in the denominator and are given the same weight as AUC I and AUC G. This is mainly a problem in cases where fasting insulin is low, which makes the Matsuda index very high. The poor agreement we found between Matsuda and MINMOD SI in the FWS data set is a consequence of a few individuals with low fasting insulin concentrations, which appear in the denominator and make the Matsuda ISI very large (not shown). The ISS model considers the fasting values in the context of fitting the entire time course and thus is much less susceptible to variability.

A similar issue arises with IGI. It is possible for glucose at 30 minutes to be close to the value at 0 minutes, which results in implausibly large IGI values. If glucose at 30 minutes is less than the value at 0 minutes, IGI is negative and again biologically implausible (36). The ISS model is not susceptible to these problems.

### Comparison with other differential equation-based fitting methods

The models most similar to the ISS model are the classical minimal model (MINMOD) (10, 37) and the oral minimal models from the Cobelli group (5, 25, 38), which also involve fitting data to differential equations. We have already discussed the good agreement between our model and MINMOD but point out here in addition that our model is not subject to the systematic underestimation of insulin sensitivity that can occur with MINMOD for individuals with high AIRg (39). As shown in that paper, the MINMOD estimate for S_I_ is inversely related to the size of AIRg, whereas the ISS model estimate for mS_I_ is insensitive to changes in AIR, modeled in synthetic data as an increase in the size of the readily releasable pool of insulin vesicles (not shown). We believe this difference stems from the sensitivity of MINMOD S_I_ to the sharp rise in insulin secretion during the first few minutes of an IM-FSIGT, which is not a major factor during an OGTT.

The method of Cobelli et al. consists of two separate models, one for S_I_ and one for BCF (5). The oral glucose minimal model is used to estimate S_I_ from an OGTT, usually with at least seven time points. Estimates of S_I_ using the oral glucose minimal model and the ISS model both correlated with clamp S_I_: the oral glucose minimal model S_I_ had a Pearson correlation of R = 0.81 in (40) and the ISS model S_I_ had a Pearson correlation of R = 0.66 in the UPMC CHP cohort. The differences in correlation coefficient between the two studies could be related to small sample size in ref 38 and/or post-hoc comparison of two very different study cohorts (middle-aged moderately obese adults vs. very obese, insulin resistant adolescents). Longer OGTTs may also produce better correlation (6): the Spearman correlation coefficient between OMM S_I_ and clamp with two-hour, six-timepoint OGTTs in the AIRS cohort of adolescent females was only R = 0.64, P < 0.0001) (41). Fitting the ISS to the same data resulted in a correlation of 0.52 (P < 0.001).

Both the oral glucose minimal model and the ISS model produced insulin sensitivity estimates that were numerically lower than clamp S_I_. The underestimation of insulin sensitivity measures derived from OGTT compared with HIEC is consistent across studies. The oral glucose minimal model S_I_ was 34% lower pairwise than clamp in that study (NS with a sample size of only 21 (40)), whereas mS_I_ was 31% lower than clamp in the UPMC CHP data set (Fig. 7A). We found that Matsuda ISI also underestimated clamp S_I_ by 20% (Fig. 7C), which was consistent with the study that introduced the Matsuda index and showed underestimation relative to clamp S_I_ (32). HOMA IS also markedly underestimated clamp S_I_. As shown in Fig. S1 C, D it was 91% smaller pairwise than clamp IS (0.17±0.01 vs. 2.1±0.1, P < 0.001). HOMA IS also markedly underestimated MINMOD S_I_: it was 72% smaller pairwise (Fig. S1 A, B). The extreme difference in agreement between HOMA IS and clamp presumably reflects the use of only fasting values in HOMA IS. This consistent underestimation of insulin sensitivity measures derived from OGTT compared with HIEC may be due to the fact that OGTT estimates reflect whole-body, not just peripheral, insulin sensitivity or to differences in the physiology evoked by the tests. However, differences in agreement between measures of insulin sensitivity may also arise due to differences in the ways individual measures are calculated and the resulting associations between an insulin sensitivity value and the reality of underlying physiology. Thus, care should be taken in comparing estimates of insulin sensitivity using different methods.

The oral C-peptide minimal model is used to estimate three quantities for BCF, phi_d, representing first-phase BCF, phi_s, representing second-phase BCF, and phi_tot, a weighted average of phi_s and phi_d. A Spearman correlation of 0.75 was reported between phi_tot and AIRg (42). The ISS measure of beta-cell function (mBCF or σ) for FWS performed similarly, with a Spearman correlation of 0.70 with AIRg (Table 4). Another study reported a Pearson correlation between phi_s (from MMT, not OGTT) and HGC second-phase BCF of R = 0.69 (43). This is comparable to the Pearson correlation of the ISS *σ* with HGC second-phase in the UPMC CHP study (R = 0.64; Fig. 8B). Thus, the performance of the ISS BCF measure is comparable to those from the C-peptide minimal model.

A methodological difference between our model and the oral glucose minimal model is that we do not include a variable for the unmeasured interstitial insulin (X in the minimal model family). We believe that X is mainly needed, along with frequent sampling, to get good fits to glucose at early time points, t < 20 min during both IVGTT using MINMOD and OGTT when using OMM. Our model estimates agree well with MINMOD and do not need X because the model does not use any time points between 0 and 30 min. Using the CU data set (20), we found that adding time points at 10 and 20 min, as used in the Oral Glucose and C-Peptide minimal models, did not improve fitting error at the remaining time points (Figs. S5 and S6). Also, the estimates of both mS_I_ and σ using five points and seven points were very highly correlated with no significant systematic disagreement (Fig. S7).

This ISS model provided good mS_I_ estimates even with three glucose and insulin time points (t = 0, 60, 120) in our three independent data sets (Figs. 9, 10, S3, S4, S8). We found that the model was still able to stratify study participants by glucose tolerance status with three time points.

Some apparent discrepancies in Fig. 11 for the UPMC CHP and CU data sets arose because the model is based on OGTT glucose and insulin, but some participants in UPMC CHP were on diabetes medications that normalized their OGTTs, and glucose tolerance for the CU participants was determined by A1c rather than glucose. In all three data sets where we compared three vs. five points, the discrepancy for mS_I_ was negligible (< 5%), albeit non-uniform in two of the data sets, whereas mBCF (σ) was underestimated on average by 15 – 25%. The underestimation of BCF means that the ISS model estimates are conservative and of value for screening for T2D risk: the model would have high sensitivity for detecting people with poor beta-cell function. Combined with low insulin sensitivity, which can be well resolved with three time points, low beta-cell function is a strong indicator of T2D risk.

The ability to estimate mS_I_ and σ precisely and accurately with reduced sampling time points (three) is advantageous in diabetes epidemiology. Large epidemiological studies, such as the Diabetes Prevention Program (44), have collected OGTTs with glucose at three timepoints (t = 0, 30, 120) and insulin at two timepoints (t = 0, 30) and applied HOMA IS, HOMA Beta and IGI. The Korean Diabetes Prevention Study used five glucose time points (0, 30, 60, 90, 120) and two insulin time points (0, 30) (45). Preliminary analysis down-sampling these data sets indicates that the ISS model can provide adequate estimates with such minimal sampling schedules (data not shown). This is another advantage of the model.

Finally, the ISS model does not require deconvolution because derivatives of sparsely sampled C-peptide and glucose time courses are not needed to estimate BCF. This simplifies the data fitting and potentially will make our model accessible to a much wider set of clinical investigators.

### Limitations

Study limitations include not having a single data set with OGTT, IM-FSIGT, and clamp measures to test and validate our findings. However, the absence of a single data set is not surprising given the complexity and burden of intravenous tests. We validated the model in both youth and women, groups at high risk for developing diabetes, but the data set was not suitable for testing reproducibility. Although we studied adolescent boys in one data set, none of the data sets included adult men. Other limitations include the increased error with high sigma and the correlation between mSI and sigma because they are derived from the same model.

### Conclusions

We have demonstrated and validated a differential equation model for estimating insulin sensitivity and beta-cell function from a five-point OGTT that strikes a balance between fidelity and pragmatic considerations. It correlated and agreed well with estimates obtained by minimal model analysis of IM-FSIGTTs from the Federal Women’s Study while requiring considerably less data. It also correlated well with estimates from HIEC and HGC in a study of obese adolescents while requiring less analytic effort and data. Performance was as good as or better than commonly used algebraic indices from OGTTs with modest additional effort. This model should be considered for use in larger studies because of its strong correlation with gold-standard measures without the participant and resource burden and with advantages of requiring fewer data points and less analytical effort.

## Supplemental Material

**Supplemental Table 1.** Contemporary Techniques for Measuring Insulin Sensitivity and Beta cell function

| Insulin sensitivity |  |  |  |  |  |
| --- | --- | --- | --- | --- | --- |
| Technique | Special resources and services for implementation | Correlation with Hyperinsulinemia euglycemic clamp | References | Correlation with FSIVGTT<sup>Insulin-Modified</sup> | References |
| FSIVGTT<sup>Insulin-Modified</sup> | Clinical Research Unit<br>Intravenous insulin administration<br>> 10 blood draws<br>3-hour test<br>Mathematical modeling | r=0.62; P<0.01 | (46)<br>11 NGT, 20 IGT, 24 T2D |  |  |
| MMT minimal model | Clinical Research Unit<br>7-10 blood draws<br>2-5-hour test<br>Mathematical modeling | r = 0.76, P<0.01 | (43)<br>16 NGT/ 1IGT | r = 0.59, P<0.01<br>r = 0.73, p<0.01 | (43)<br>16 NGT/ 1IGT<br>(25) 22 NGT |
| OGTT glucose minimal model | Clinical Research Unit<br>7-10 blood draws<br>2-3-hour test<br>Mathematical modeling | r=0.81, P<0.001 | (40)<br>10 NGT, 11 IGT |  |  |
| OGTT Matsuda index | Clinical Research Unit<br>3-5 blood draws<br>2-hour test<br>Simple algebraic equation | r>0.73; P<0.01 | (32)<br>62 NGT, 31 IGT, 60 T2D | r=0.68, P<0.05 | (47)<br>20 NGT |
| HOMA-IR | Single fasting blood draw<br>Simple algebraic equation | r=-0.88, P<0.01 | (33)<br>12 NGT, 11 T2D | r=-0.56, P<0.05 | (47)<br>20 NGT |
| Beta cell function |  |  |  |  |  |
| Technique |  | Hyperglycemic clamp 1<sup>st</sup> phase/ 2<sup>nd</sup> phase | References | AIRg (scales are different) | References |
| FSIGT - AIRg | Clinical Research Unit<br>Intravenous glucose administration<br>4-10 blood draws<br>10 minutes<br>Simple algebraic equation | r = 0.96, P<0.0001<br>r=0.75, P<0.005 | (43)<br>16 NGT/ 1 IGT<br>(48)<br>14 NGT |  |  |
| OGTT insulinogenic index | Clinical Research Unit<br>3 blood draws<br>0.5 - 2-hour test<br>Simple algebraic equation | r=0.86, P<0.05 (1<sup>st</sup> phase)<br>r=0.80, P<0.05 (2<sup>nd</sup> phase) | (49) 31 NGT | r=0.52, p=0.005<br>r=0.61, P<0.001<br>r=0.49, P<0.01 | (50)<br>(31)<br>85 NGT, 23 IGT<br>(30)<br>469 NGT |
| MMT C-peptide minimal model | Clinical Research Unit<br>7-10 blood draws<br>Mathematical modeling | r=0.288, P=0.2617 (1<sup>st</sup> phase)<br>r = 0.686, P<0.01 (2<sup>nd</sup> phase) | (43)<br>16 NGT/ 1 IGT | r=0.282, P=0.2731 | (43)<br>16 NGT/ 1 IGT |
| OGTT insulin AUC/ glucose AUC | Clinical Research Unit<br>2-5 blood draws<br>Simple algebraic equation |  |  | r=0.32, p=0.08 | (50)<br>40 NGT |
| HOMA-Beta | Single fasting blood draw<br>Simple algebraic equation | r=0.61, P<0.01<br>r=0.69, P<0.05 (1<sup>st</sup> phase)<br>r=0.72, P<0.05 (2<sup>nd</sup> phase) | (33)<br>10 NGT/ 10T2D<br>(49)<br>31 NGT | r=0.64, P<0.05<br>r=0.58, p<0.01 | (33)<br>6 NGT/ 6T2D<br>(51)<br>712 NGT, 353 IGT,<br>315 T2D |

Homeostasis Model Assessment Insulin Resistance:HOMA-IR; Oral Glucose tolerance test OGTT; frequently sampled intravenous glucose tolerance test FSIVGTT; AIRg: acute insulin response to glucose; AUC: area under curve; NGT: normal glucose tolerant;T2D: type 2 diabetes mellitus. r=correlation coefficient

Supplemental Figures S1 – S8

Supplemental Computer Code

Supplemental Methods

Supplemental Equations

All at doi: https://doi.org/10.6084/m9.figshare.23535612

## Acknowledgments

The work of ASS, JH, STC, PC, and MS was supported by the Intramural Research Program of the National Institutes of Health (NIDDK). JH was also supported by a dkNET New Investigator Pilot Program in Bioinformatics, NIDDK, and the Brain Pool Program of the National Research Foundation of Korea (grant # 2022H1D3A2A01063552). MS was also supported by the National Science Foundation Graduate Research Fellowship Program under Grant No. DGE 1840340. Any opinions, findings, and conclusions or recommendations expressed in this material are those of the authors and do not necessarily reflect the views of the National Science Foundation. The work of SA was supported by NIH R01 HD27503, NIH K24 HD/DK01357, and NIH UL1TR001857 and RR024153. The work of MGC was supported by NCATS CTSI UL1 TR002535, BIRCWH K12HD057022, NIDDK K23DK107871, NIDDK R01DK120612, Doris Duke Foundation CDSA, and a Boettcher Webb-Waring award. The work of CDB was supported by NSF DMS 1853511. We thank Raibatak Das (Applied BioMath, Inc). for making his lecture notes on bootstrapping regression available. We thank Katrina Diesche (NIDDK) for testing the usability of the model and providing valuable feedback.

## References

1. Ishimwe MCS, Wentzel A, Shoup EM, Osei-Tutu NH, Hormenu T, Patterson AC, Bagheri H, DuBose CW, Mabundo LS, Ha J, Sherman A, and Sumner AE. Beta-cell failure rather than insulin resistance is the major cause of abnormal glucose tolerance in Africans: insight from the Africans in America study. BMJ Open Diabetes Res Care 9: 2021.

2. Esser N, Utzschneider KM, and Kahn SE. On the causal relationships between hyperinsulinaemia, insulin resistance, obesity and dysglycaemia in type 2 diabetes: Reply to Johnson JD [letter]. Diabetologia 64: 2345–2347, 2021.

3. Johnson JD. On the causal relationships between hyperinsulinaemia, insulin resistance, obesity and dysglycaemia in type 2 diabetes. Diabetologia 64: 2138–2146, 2021.

4. Ohn JH, Kwak SH, Cho YM, Lim S, Jang HC, Park KS, and Cho NH. 10-year trajectory of beta-cell function and insulin sensitivity in the development of type 2 diabetes: a community-based prospective cohort study. Lancet Diabetes Endocrinol 4: 27–34, 2016.

5. Cobelli C, Dalla Man C, Toffolo G, Basu R, Vella A, and Rizza R. The oral minimal model method. Diabetes 63: 1203–1213, 2014.

6. Bartlette K, Carreau AM, Xie D, Garcia-Reyes Y, Rahat H, Pyle L, Nadeau KJ, Cree-Green M, and Diniz Behn C. Oral minimal model-based estimates of insulin sensitivity in obese youth depend on oral glucose tolerance test protocol duration. Metabol Open 9: 100078, 2021.

7. Chung ST, Ha J, Onuzuruike AU, Kasturi K, Galvan-De La Cruz M, Bingham BA, Baker RL, Utumatwishima JN, Mabundo LS, Ricks M, Sherman AS, and Sumner AE. Time to glucose peak during an oral glucose tolerance test identifies prediabetes risk. Clin Endocrinol (Oxf) 87: 484–491, 2017.

8. Chung ST, Galvan-De La Cruz M, Aldana PC, Mabundo LS, DuBose CW, Onuzuruike AU, Walter M, Gharib AM, Courville AB, Sherman AS, and Sumner AE. Postprandial Insulin Response and Clearance Among Black and White Women: The Federal Women’s Study. J Clin Endocrinol Metab 104: 181–192, 2019.

9. Chung ST, Cravalho CKL, Meyers AG, Courville AB, Yang S, Matthan NR, Mabundo L, Sampson M, Ouwerkerk R, Gharib AM, Lichtenstein AH, Remaley AT, and Sumner AE. Triglyceride Paradox Is Related to Lipoprotein Size, Visceral Adiposity and Stearoyl-CoA Desaturase Activity in Black Versus White Women. Circ Res 126: 94–108, 2020.

10. Boston RC, Stefanovski D, Moate PJ, Sumner AE, Watanabe RM, and Bergman RN. MINMOD Millennium: a computer program to calculate glucose effectiveness and insulin sensitivity from the frequently sampled intravenous glucose tolerance test. Diabetes Technol Ther 5: 1003–1015, 2003.

11. Sjaarda LA, Michaliszyn SF, Lee S, Tfayli H, Bacha F, Farchoukh L, and Arslanian SA. HbA(1c) diagnostic categories and beta-cell function relative to insulin sensitivity in overweight/obese adolescents. Diabetes Care 35: 2559–2563, 2012.

12. Burns SF, Bacha F, Lee SJ, Tfayli H, Gungor N, and Arslanian SA. Declining beta-cell function relative to insulin sensitivity with escalating OGTT 2-h glucose concentrations in the nondiabetic through the diabetic range in overweight youth. Diabetes Care 34: 2033–2040, 2011.

13. Kim JY, Tfayli H, Bacha F, Lee S, Gebara N, and Arslanian S. beta-cell impairment and clinically meaningful alterations in glycemia in obese youth across the glucose tolerance spectrum. Metabolism 112: 154346, 2020.

14. Sjaarda LG, Bacha F, Lee S, Tfayli H, Andreatta E, and Arslanian S. Oral disposition index in obese youth from normal to prediabetes to diabetes: relationship to clamp disposition index. J Pediatr 161: 51–57, 2012.

15. Kim JY, Michaliszyn SF, Nasr A, Lee S, Tfayli H, Hannon T, Hughan KS, Bacha F, and Arslanian S. The Shape of the Glucose Response Curve During an Oral Glucose Tolerance Test Heralds Biomarkers of Type 2 Diabetes Risk in Obese Youth. Diabetes Care 39: 1431–1439, 2016.

16. Kim JY, Bacha F, Tfayli H, Michaliszyn SF, Yousuf S, and Arslanian S. Adipose Tissue Insulin Resistance in Youth on the Spectrum From Normal Weight to Obese and From Normal Glucose Tolerance to Impaired Glucose Tolerance to Type 2 Diabetes. Diabetes Care 42: 265–272, 2019.

17. George L, Bacha F, Lee S, Tfayli H, Andreatta E, and Arslanian S. Surrogate estimates of insulin sensitivity in obese youth along the spectrum of glucose tolerance from normal to prediabetes to diabetes. J Clin Endocrinol Metab 96: 2136–2145, 2011.

18. Bacha F, Gungor N, Lee S, and Arslanian SA. In vivo insulin sensitivity and secretion in obese youth: what are the differences between normal glucose tolerance, impaired glucose tolerance, and type 2 diabetes? Diabetes Care 32: 100–105, 2009.

19. Teede HJ, Misso ML, Costello MF, Dokras A, Laven J, Moran L, Piltonen T, Norman RJ, and International PN. Recommendations from the international evidence-based guideline for the assessment and management of polycystic ovary syndrome. Hum Reprod 33: 1602–1618, 2018.

20. Carreau AM, Jin ES, Garcia-Reyes Y, Rahat H, Nadeau KJ, Malloy CR, and Cree-Green M. A simple method to monitor hepatic gluconeogenesis and triglyceride synthesis following oral sugar tolerance test in obese adolescents. Am J Physiol Regul Integr Comp Physiol 317: R134–R142, 2019.

21. Cree-Green M, Bergman BC, Coe GV, Newnes L, Baumgartner AD, Bacon S, Sherzinger A, Pyle L, and Nadeau KJ. Hepatic Steatosis is Common in Adolescents with Obesity and PCOS and Relates to De Novo Lipogenesis but not Insulin Resistance. Obesity (Silver Spring) 24: 2399–2406, 2016.

22. DiMeglio LA, Acerini CL, Codner E, Craig ME, Hofer SE, Pillay K, and Maahs DM. ISPAD Clinical Practice Consensus Guidelines 2018: Glycemic control targets and glucose monitoring for children, adolescents, and young adults with diabetes. Pediatr Diabetes 19 Suppl 27: 105–114, 2018.

23. Ha J, Satin LS, and Sherman AS. A Mathematical Model of the Pathogenesis, Prevention, and Reversal of Type 2 Diabetes. Endocrinology 157: 624–635, 2016.

24. Ha J, and Sherman A. Type 2 diabetes: one disease, many pathways. Am J Physiol Endocrinol Metab 319: E410–E426, 2020.

25. Dalla Man C, Caumo A, and Cobelli C. The oral glucose minimal model: estimation of insulin sensitivity from a meal test. IEEE Trans Biomed Eng 49: 419–429, 2002.

26. Topp B, Promislow K, deVries G, Miura RM, and Finegood DT. A model of beta-cell mass, insulin, and glucose kinetics: pathways to diabetes. J Theor Biol 206: 605–619, 2000.

27. Chen YD, Wang S, and Sherman A. Identifying the targets of the amplifying pathway for insulin secretion in pancreatic beta-cells by kinetic modeling of granule exocytosis. Biophys J 95: 2226–2241, 2008.

28. Grodsky GM. A threshold distribution hypothesis for packet storage of insulin and its mathematical modeling. J Clin Invest 51: 2047–2059, 1972.

29. Efron B, and Tibshirani RJ. An Introduction to the Bootstrap. Boca Raton: Chapman & Hall/CRC, 1993, p. 456.

30. Hanson RL, Pratley RE, Bogardus C, Narayan KM, Roumain JM, Imperatore G, Fagot-Campagna A, Pettitt DJ, Bennett PH, and Knowler WC. Evaluation of simple indices of insulin sensitivity and insulin secretion for use in epidemiologic studies. Am J Epidemiol 151: 190–198, 2000.

31. Phillips DI, Clark PM, Hales CN, and Osmond C. Understanding oral glucose tolerance: comparison of glucose or insulin measurements during the oral glucose tolerance test with specific measurements of insulin resistance and insulin secretion. Diabet Med 11: 286–292, 1994.

32. Matsuda M, and DeFronzo RA. Insulin sensitivity indices obtained from oral glucose tolerance testing: comparison with the euglycemic insulin clamp. Diabetes Care 22: 1462–1470, 1999.

33. Matthews DR, Hosker JP, Rudenski AS, Naylor BA, Treacher DF, and Turner RC. Homeostasis model assessment: insulin resistance and beta-cell function from fasting plasma glucose and insulin concentrations in man. Diabetologia 28: 412–419, 1985.

34. Altman DG, and Bland JM. Measurement in Medicine: The Analysis of Method Comparison Studies. Journal of the Royal Statistical Society: Series D 32: 307–317, 1983.

35. Consortium R. Metabolic Contrasts Between Youth and Adults With Impaired Glucose Tolerance or Recently Diagnosed Type 2 Diabetes: I. Observations Using the Hyperglycemic Clamp. Diabetes Care 41: 1696–1706, 2018.

36. Faulenbach MV, Wright LA, Lorenzo C, Utzschneider KM, Goedecke JH, Fujimoto WY, Boyko EJ, McNeely MJ, Leonetti DL, Haffner SM, Kahn SE, and American Diabetes Association GSG. Impact of differences in glucose tolerance on the prevalence of a negative insulinogenic index. J Diabetes Complications 27: 158–161, 2013.

37. Bergman RN, Ider YZ, Bowden CR, and Cobelli C. Quantitative estimation of insulin sensitivity. Am J Physiol 236: E667–677, 1979.

38. Breda E, Cavaghan MK, Toffolo G, Polonsky KS, and Cobelli C. Oral glucose tolerance test minimal model indexes of beta-cell function and insulin sensitivity. Diabetes 50: 150–158, 2001.

39. Ha J, Muniyappa R, Sherman AS, and Quon MJ. When MINMOD Artifactually Interprets Strong Insulin Secretion as Weak Insulin Action. Front Physiol 12: 601894, 2021.

40. Dalla Man C, Yarasheski KE, Caumo A, Robertson H, Toffolo G, Polonsky KS, and Cobelli C. Insulin sensitivity by oral glucose minimal models: validation against clamp. Am J Physiol Endocrinol Metab 289: E954–959, 2005.

41. Carreau AM, Xie D, Garcia-Reyes Y, Rahat H, Bartlette K, Behn CD, Pyle L, Nadeau KJ, and Cree-Green M. Good agreement between hyperinsulinemic-euglycemic clamp and 2 hours oral minimal model assessed insulin sensitivity in adolescents. Pediatr Diabetes 21: 1159–1168, 2020.

42. Shankar SS, Vella A, Raymond RH, Staten MA, Calle RA, Bergman RN, Cao C, Chen D, Cobelli C, Dalla Man C, Deeg M, Dong JQ, Lee DS, Polidori D, Robertson RP, Ruetten H, Stefanovski D, Vassileva MT, Weir GC, Fryburg DA, and Foundation for the National Institutes of Health beta-Cell Project T. Standardized Mixed-Meal Tolerance and Arginine Stimulation Tests Provide Reproducible and Complementary Measures of beta-Cell Function: Results From the Foundation for the National Institutes of Health Biomarkers Consortium Investigative Series. Diabetes Care 39: 1602–1613, 2016.

43. Steil GM, Hwu CM, Janowski R, Hariri F, Jinagouda S, Darwin C, Tadros S, Rebrin K, and Saad MF. Evaluation of insulin sensitivity and beta-cell function indexes obtained from minimal model analysis of a meal tolerance test. Diabetes 53: 1201–1207, 2004.

44. Kitabchi AE, Temprosa M, Knowler WC, Kahn SE, Fowler SE, Haffner SM, Andres R, Saudek C, Edelstein SL, Arakaki R, Murphy MB, Shamoon H, and Diabetes Prevention Program Research G. Role of insulin secretion and sensitivity in the evolution of type 2 diabetes in the diabetes prevention program: effects of lifestyle intervention and metformin. Diabetes 54: 2404–2414, 2005.

45. Rhee SY, Chon S, Ahn KJ, Woo JT, and Korean Diabetes Prevention Study I. Hospital-Based Korean Diabetes Prevention Study: A Prospective, Multi-Center, Randomized, Open-Label Controlled Study. Diabetes Metab J 43: 49–58, 2019.

46. Saad MF, Anderson RL, Laws A, Watanabe RM, Kades WW, Chen YD, Sands RE, Pei D, Savage PJ, and Bergman RN. A comparison between the minimal model and the glucose clamp in the assessment of insulin sensitivity across the spectrum of glucose tolerance. Insulin Resistance Atherosclerosis Study. Diabetes 43: 1114–1121, 1994.

47. Henderson M, Rabasa-Lhoret R, Bastard JP, Chiasson JL, Baillargeon JP, Hanley JA, and Lambert M. Measuring insulin sensitivity in youth: How do the different indices compare with the gold-standard method? Diabetes Metab 37: 72–78, 2011.

48. Korytkowski MT, Berga SL, and Horwitz MJ. Comparison of the minimal model and the hyperglycemic clamp for measuring insulin sensitivity and acute insulin response to glucose. Metabolism: clinical and experimental 44: 1121–1125, 1995.

49. Uwaifo GI, Fallon EM, Chin J, Elberg J, Parikh SJ, and Yanovski JA. Indices of insulin action, disposal, and secretion derived from fasting samples and clamps in normal glucose-tolerant black and white children. Diabetes Care 25: 2081–2087, 2002.

50. Powe CE, Locascio JJ, Gordesky LH, Florez JC, and Catalano PM. Oral Glucose Tolerance Test-based Measures of Insulin Secretory Response in Pregnancy. J Clin Endocrinol Metab 107: e1871–e1878, 2022.

51. Festa A, Williams K, Hanley AJ, and Haffner SM. Beta-cell dysfunction in subjects with impaired glucose tolerance and early type 2 diabetes: comparison of surrogate markers with first-phase insulin secretion from an intravenous glucose tolerance test. Diabetes 57: 1638–1644, 2008.

